# Regulation and Function of a Polarly Localized Lignin Barrier in the Exodermis

**DOI:** 10.1101/2022.10.20.513117

**Authors:** Concepcion Manzano, Kevin W. Morimoto, Lidor Shaar-Moshe, G. Alex Mason, Alex Cantó-Pastor, Mona Gouran, Damien De Bellis, Robertas Ursache, Kaisa Kajala, Neelima Sinha, Julia Bailey-Serres, Niko Geldner, J Carlos del Pozo, Siobhan M. Brady

## Abstract

Multicellular organisms control interactions with their environment through the development of specialized barriers in specific cell types. A conserved barrier in plant roots is the endodermal Casparian strip (CS). The CS is made of polymerized lignin and forms a ring-like structure that seals the apoplastic space between the endodermal cells. Most angiosperms also have another root cell type, the exodermis, that is reported to form a barrier. Our understanding of exodermal developmental and molecular regulation, as well as function, is limited as this cell type is absent from the model species *Arabidopsis thaliana*. Using tomato (*Solanum lycopersicum*) as a model system we demonstrate that in this species, the exodermis does not form a CS. Instead, it forms a polar lignin cap with an equivalent barrier function to the endodermal CS. We demonstrate that although endodermal regulators are conserved between Arabidopsis and tomato, exodermal differentiation occurs by a distinct regulatory pathway involving the *SlSCZ* and *SlEXO1* transcription factors. Although the exodermis and endodermis both produce barriers that restrict mineral ion uptake, they have unique and overlapping roles in their selectivity. Whether conservation and similarities between the endodermis and exodermis exist in other species remains to be determined. Nonetheless, in tomato, these distinct lignin structures have a convergent function with different genetic regulations.

## INTRODUCTION

Plant roots anchor the plant and absorb water and nutrients. The control of external mineral ions and water entry is essential for plant survival. In vascular plants, the endodermis is the innermost root cell layer that controls apoplastic movement to and from the plant vasculature by the formation of a barrier that seals the intercellular (apoplastic) space. A first step in endodermal differentiation is the formation of the Casparian strip (CS) which controls the free diffusion of solutes from the soil and prevents the backflow of ions that have been accumulated in the stele (Enstone, Peterson, and Ma 2002).

The developmental framework of endodermis differentiation has been well elucidated in *Arabidopsis thaliana* (Alassimone, Naseer, and Geldner 2010; Roppolo et al. 2011; Hosmani et al. 2013; Y. Lee et al. 2013; Doblas et al. 2017; Nakayama et al. 2017). Specification of endodermis identity occurs within the cortex-endodermis stem cell niche (Helariutta et al. 2000) via the non-cell autonomous action of the SHORT-ROOT (SHR) transcription factor. Its subsequent transcriptional cascade coordinates the expression of factors that synthesize, deposit, and position the CS (Benfey et al. 1993; Gallagher et al. 2004; Helariutta et al. 2000; Nakajima et al. 2001).

In the primary differentiation stage, the CS is formed as a localized impregnation of the primary cell wall with polymerized lignin forming a ring around the central axis of the cell (Naseer et al. 2012). In the more conservative definition of a CS, this lignin is deposited in a very precise and localized central position within the anticlinal cell walls (von Guttenberg 1968). Hallmarks of this precise positioning include tight membrane adhesion and the action and localization of protein domains comprised of CASPARIAN STRIP DOMAIN PROTEIN (CASP)/CASP-like and the ENHANCED SUBERIN 1 (ESB1) proteins (Alassimone, Naseer, and Geldner 2010; Roppolo et al. 2011; Hosmani et al. 2013; Y. Lee et al. 2013; Rojas-Murcia et al., 2020). The transcription factor AtMYB36 positively regulates the expression of the *AtCASP1*, *AtPER64*, and *AtESB1* genes that are necessary to define CS positioning as well as its polymerization. Mutation of *AtMYB36* (*atmyb36*) results in an absent CS (Kamiya et al. 2015; Liberman et al. 2015), ectopic lignin deposition in endodermal cell corners as well as disruption of CS barrier function (Kamiya et al. 2015). The *SCHENGEN3/SCHENGEN1/CASPARIAN STRIP INTEGRITY FACTOR2(SGN3/SGN1/CIF2*) pathway acts as an elegant surveillance system to perceive defects in CS integrity. If such defects occur, this pathway activates compensatory lignification and suberization (Doblas et al. 2017; Nakayama et al. 2017). Compensatory corner lignification in the *athmyb36* mutant is mediated by the SGN3/SGN1/CIF pathway as the double mutant *atmyb36atsgn3* presents a complete loss in endodermis lignification (Reyt et al. 2021). Both the *atmyb36* and the double mutant *atmyb36atsgn3* have a drastic impact on ion homeostasis and growth (Kamiya et al., 2015; Reyt et al., 2021) proving the important function of endodermal CS in whole plant growth. The second differentiation stage is marked by the deposition of suberin lamellae that eventually coat the entire endodermal cell wall surface (Barberon et al. 2016; Geldner 2013).

Many plant species, but not Arabidopsis, also contain an additional cell type that has barrier function, the exodermis or the hypodermis, herein referred to as the exodermis. Located underneath the epidermis, the exodermis is present in more than 90% of 200 angiosperms previously examined (Perumalla, Peterson, and Enstone 1990; Peterson and Perumalla 1990; Perumalla, Chmielewski, and Peterson 1990). Based on the staining technology available at this time, it was shown that exodermal cell walls contain lignin- or suberin-related compounds, in a variety of deposition patterns, many of which included a broad band along the radial and anticlinal cell walls. The methods used did not allow a straightforward differentiation between lignin and suberin within these walls. At the time, a Casparian strip was thought to be composed of both suberin and lignin and the authors therefore concluded that the exodermis in these species possesses a CS. More recently, however, a CS has been conclusively demonstrated to be composed solely of polymerized lignin, which is necessary for its function as an apoplastic barrier. Unanswered questions include whether the exodermis contains lignin or suberin, its precise subcellular location and which genes control exodermal differentiation.

Cell population translatome profiles and gene ontology enrichment in domesticated tomato, *Solanum lycopersicum*, cv. M82 allowed us to hypothesize that the tomato exodermis is both lignified and suberized, which we subsequently validated experimentally (Kajala et al. 2021). The exodermis in tomato has two stages of differentiation: first, a polar lignin cap is deposited and localized on the epidermal face of exodermal cells, followed by suberization surrounding the entire cell surface. Here, we demonstrate that in tomato, the exodermal barrier is not a CS, but the polar lignin cap nevertheless functions as an apoplastic diffusion barrier and describes the developmental timeline by which this structure forms. We further demonstrate that while regulation of endodermal CS differentiation is generally conserved between Arabidopsis and tomato, the regulatory pathway for the polar lignin cap in exodermal differentiation is genetically distinct. A moderate throughput CRISPR-Cas9 screen identified two transcription factors, *SlEXO1* and *SlSCHIZORIZA (SlSCZ*) which collectively restrict the deposition of the polar lignin cap to the exodermis and *SlSCZ* additionally regulates the polarity of the lignin cap. Phenotypic analysis of these mutants shows that the exodermal polar cap selectively restricts mineral ion uptake with similar and unique properties compared to those of the endodermal CS. Both genes coordinate multiple downstream regulatory pathways to synthesize the lignin cap including genes involved in cell wall remodeling, lignin biosynthesis, and phenylpropanoid regulation. Our findings reveal for the first time that exodermal differentiation is distinct from endodermal specification and differentiation and that its polar cap acts as a first line apoplastic barrier, with an equivalent apoplastic barrier function.

## RESULTS

### Exodermal lignification comprises a polarly localized lignin cap that acts as an apoplastic barrier

The CS in the *Arabidopsis* endodermis is composed of polymerized lignin impregnated in the cell wall, forming a narrow belt surrounding the longitudinal, central axis of the cell. Endodermal lignification occurs in the early root maturation zone (Naseer et al. 2012). In tomato, (*Solanum lycopersicum*) the exodermis forms a polarized lignin cap on the epidermal facing exodermal cell surface (**Figure 1A**). The temporal nature of exodermal lignin deposition was characterized in five-day-old seedlings using histochemical staining with Basic Fuchsin (lignin) and Calcofluor White (cellulose) to counterstain the cell walls (**Figure 1A**). These data were complemented with transmission electron microscopy (TEM) combined with potassium permanganate staining to visualize electron-dense polymerized lignin (**Fig 1C, D, Supplemental Figure 3**) (Hepler, Fosket, and Newcomb 1970). Exodermal lignification occurs after endodermal CS deposition in the early maturation zone. First, lignification occasionally occurs at exodermal epidermal cell corners, at approximately 0.4 cm from the root tip which we term Stage 1 of exodermal lignification (**Figure 1B-D**). In Stage 2 of exodermal differentiation, at around 1 cm from the root tip lignin increases at exodermal cell corners and becomes deposited all along the epidermal face of exodermal cells (**Figure 1B**). Stage 3 of exodermal differentiation occurs at around 2 cm from the root tip, with lignin content increasing and spreading along the anticlinal cell walls towards the first inner cortical cell layer. This stage is maintained from 2 to 8 cm from the root tip, near the root-hypocotyl junction where Stage 4 of exodermal differentiation comprises maximal lignification of approximately ⅔ of the anticlinal cell wall (**Figure 1B**). Polymerized lignin levels additionally increase at the epidermal face (**Figure 1B**). This polar lignin cap in the exodermis in Solanum lycopersicum is also conserved in all Solanaceous species examined and *Nicotiana benthamiana* (**Supplemental Figure 1**).

**Figure 1.**
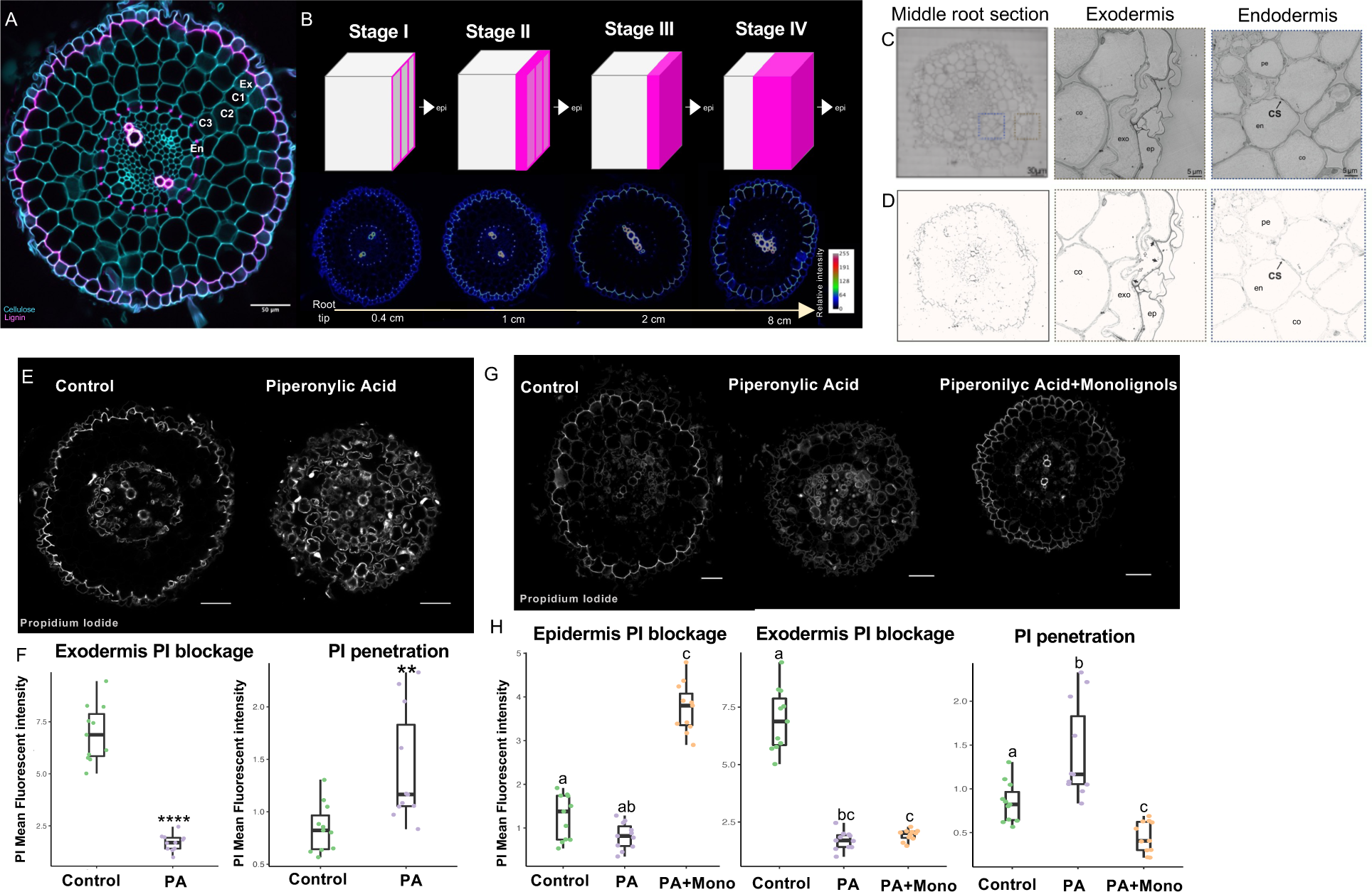
Exodermal lignin is polarized and serves as an apoplastic barrier. **A.** Tomato root cross-section stained with basic fuchsin (pink) and calcofluor (blue) for lignin and cellulose respectively. Ex = exodermis, C1 = cortex 1, C2 = cortex 2, C3 = cortex 3, En= endodermis. **B.** Model for exodermis lignin deposition. Upper part: each cube represents an exodermis cell. The epidermis face is situated on the right. Pink color represents lignin. Bottom part: root cross sections for each stage stained with basic fuchsin. The sections were taken at 0.4cm, 1cm, 2cm and 8cm from the root tip. **C.** Transmission electron microscopy (TEM) microscopy of a tomato root section taken from the middle of the root, stained with potassium permanganate (KMnO_4_). Left panel= Middle root section. Middle panel: Exodermis. Magnified grey square from left panel. Right panel: Endodermis. Magnified blue square from left panel. ep = epidermis, ex = exodermis, co = cortex, en = endodermis, xy = xylem. **D**. Same images as in C with an adapted contrast to highlight lignin deposition. **E.** Tomato root sections from control and treated plants with Piperonylic Acid (PA) for 24 hours. The plants were next incubated in the apoplastic tracer Propidium Iodide (PI) for 30 minutes. **F.** Quantification of PI blockage at the exodermal lignin cap and penetration into cortex cells in control and PA-treated plants (200 μM) for 24 hours and incubated with PI for 30 minutes (p-value for exodermis PI blockage 9.5e11 and for PI penetration 0.046 n=11. *= statistical significance as determined by ANOVA). **G.** Tomato root cross-sections from control, PA-treated plants (200 μM) and PA-treated plants with monolignols (200 μM and 20 μM respectively) for 24 hours. The plants were later incubated in the apoplastic tracer PI for 30 minutes. **H.** Quantification of PI blockage at the epidermis, the exodermal lignin cap and penetration into cortex cells in control and PA-treated plants for 24 hours followed by incubation with PI for 30 minutes (n=11; statistical significance was determined by ANOVA with a post-hoc Tukey HSD test).

The formation of an apoplastic barrier is a hallmark of CS function. To test if the polar exodermal lignin cap has an equivalent function, we used propidium iodide (PI) as an apoplastic tracer (Naseer et al. 2012; Wang et al. 2019). PI apoplastic transport was still blocked by the exodermal cap after 30 minutes of incubation as indicated by the lack of PI penetrance to inner cortical cell layers (**Figure 1E-F**). Vascular PI presence was due to PI absorption within the meristem, where an exodermal lignin cap was not present, and led to subsequent transport through the xylem. Piperonylic acid (PA), a monolignol biosynthetic inhibitor, was used to further demonstrate that exodermal barrier function is dependent upon lignin biosynthesis (Schalk et al. 1998; Naseer et al. 2012). Twenty-four hours of 200 with μM PA exposure does not interfere with root growth but does disrupt root lignin biosynthesis within the endodermis and exodermis (**Supplemental Figure 2A-B**). In PA-treated plants, PI staining of cortical cells was observed after exposure (**Figure 1E**). In addition, the PI signal in the exodermal lignin cap was more intense in control relative to PA-treated plants (**Figure 1F**). The dependence of exodermal barrier function on lignin biosynthesis was further determined by complementation of the PA inhibitor-induced defects by the addition of two components of angiosperm lignin, the monolignols coniferyl-, and sinapyl-alcohol (20 with μM each), as previously used to demonstrate CS barrier function in Arabidopsis endodermis (Naseer et al. 2012). Treatment with both PA and monolignols restored lignin levels in the exodermis, increased lignin levels within the epidermis and blocked PI transport at the epidermis (**Figure 1G-H**).

### Exodermal and endodermal differentiation are genetically distinct processes

Although structurally distinct, the tomato exodermis forms an apoplastic barrier with a similar function to the endodermal CS. Therefore, one working hypothesis is that genes that regulate endodermal differentiation also regulate exodermal differentiation. Alternatively, a distinct set of genes regulate differentiation of these cell types. Recent evidence in maize demonstrates that mutations in two of the three *ZmSHR* paralogs result in a reduction of cortex layer number and likely cortex specialization (Ortiz-Ramírez et al. 2021). The tomato exodermal layer results from asymmetric divisions of the cortex-endodermal initial at the stem cell niche (Ron et al. 2013). We therefore tested whether *SlSHR* regulates formation of the exodermal polar lignin cap. *SlSHR* (Solyc02g092370) transcript is produced in the stele, its protein is found in the endodermis, and *SlSHR* controls transcription of its downstream target *SlSCR* (Solyc10g074680) in *Rhizobium rhizogenes*-transformed roots (Hairy roots) (Ron et al. 2014). Both *R. rhizogenes* and *A. tumefaciens*-generated *SlSHR* CRISPR-Cas9 mutant alleles have defects in endodermal specification and differentiation, including the absence of the endodermal layer as well as the CS (**Figure 2A**, **Supplementary Figure 4A-B**). In parts of the root where other cell types have already differentiated (exodermal lignin cap deposition and xylem secondary cell wall thickening), the mutant layer that surrounds the vasculature has no CS or ectopic lignification and more mature cells in this layer deposit ectopic lignin (**Figure 2A**). Both *slshr-1* and *slshr-2* alleles additionally have asymmetric radial patterning in the ground tissue layers and the vascular cylinder (**Figure 2B**, **Supplementary Figure 4C-D**). Both alleles also develop shorter roots (**Figure 2B**). Phenotypes corresponding to a missing endodermal layer and a properly positioned and polymerized CS demonstrate that *SlSHR* function is largely conserved between *Arabidopsis* and tomato for endodermal patterning and differentiation. A deviation from the *Arabidopsis* mutant phenotype is the radial asymmetry in the cortex cell layers (including the exodermis), and aberrant lignin deposition in the mutant layer (**Figure 2A-B**). However, all *slshr* alleles had a wild type exodermal lignin cap (**Figure 2A**).

**Figure 2.**
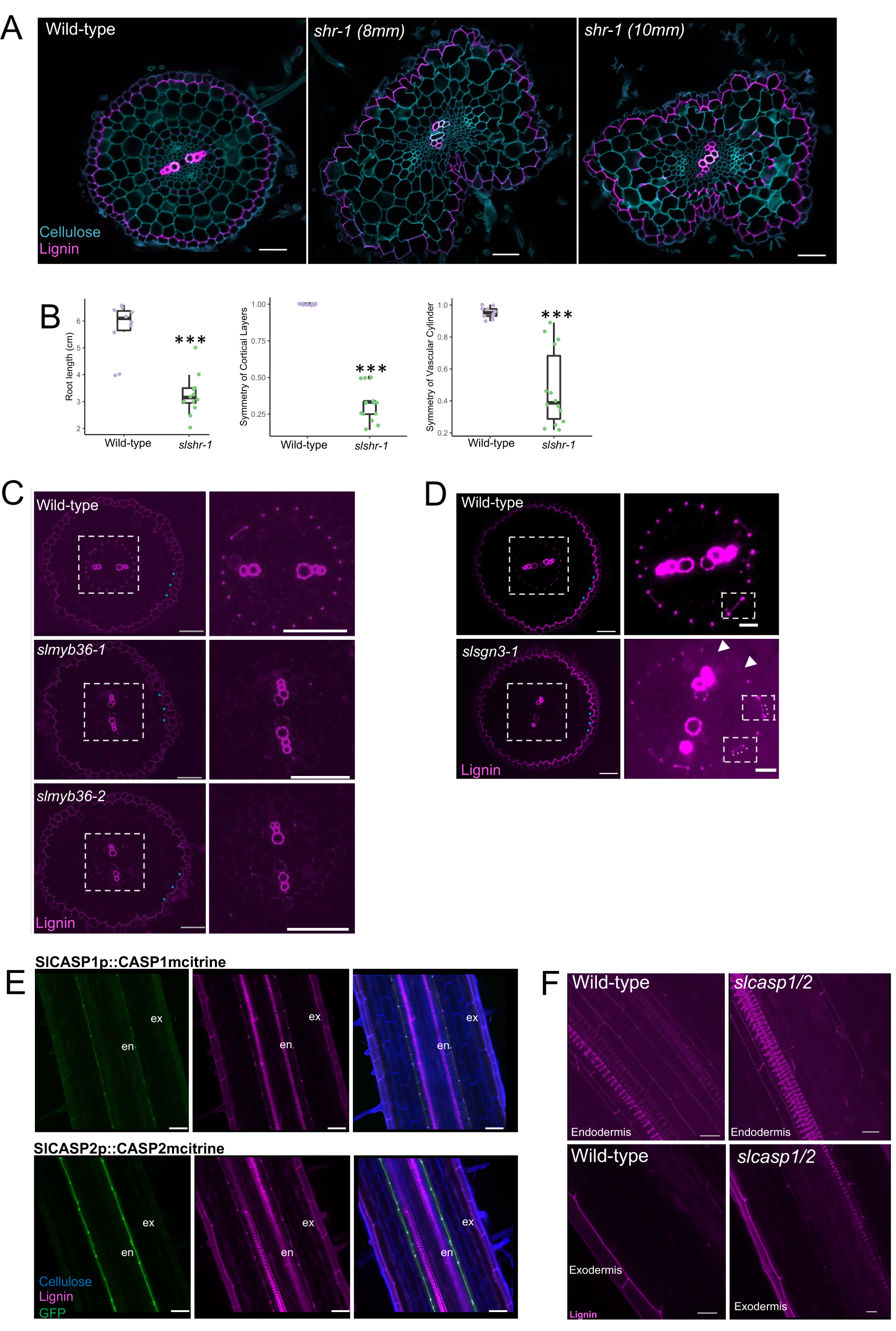
Known endodermal developmental regulators do not control exodermal differentiation. **A.** Root sections of four-day-old plants. In wild type, the endodermal layer has a classical CS composed of polymerized lignin, while the exodermis has a polar lignin cap. Basic fuchsin (pink) is used to stain lignin, while calcofluor is used to stain the cell walls (blue). The *slshr-1* mutant (middle panel) has a missing CS earlier in root development (8mm from root tip) while ectopic lignin deposition is observed in cells that surround the vasculature at 10mm from the root tip (right panel). Scale Bar = 50 with μm. **B.** Left panel: root length of four-day-old wild type and the *slshr-1* mutant (p-value=1.26e06, ANOVA). Middle panel: Symmetry of the cortical layers (including the exodermis), in radial cross-section, was calculated as the minimum number of cortex layers observed divided by the maximum number of cortex layers in wild type and *slshr-1* mutant roots (p-value=9.85e15, ANOVA). Right panel: Symmetry of the vascular cylinder was calculated as the minimum distance across the center of the vascular cylinder divided by the maximum distance across the center of the vascular cylinder (p-value=2.72e06, ANOVA) (Signif. codes: 0 ‘***’ 0.001 ‘**’ 0.01 ‘*’ 0.05) **C.** Two independent mutant alleles of *SlMYB36* (*slmyb36-1* and *slmyb36-2*) have an endodermis layer with no CS, but with a wild-type exodermal polar lignin cap. Left panels = whole root image counterstained with fuchsin, blue asterisks show the polar lignin cap in the exodermis is not affected; right panels = magnified image of vascular cylinder and endodermis layer. Scale Bar = 50 with μm **D.** Mutation in *SlSGN3* (*slsgn3-1*) results in an interrupted non-continuous CS in the radial axis. Left panels = whole root image counterstained with fuchsin, blue asterisks show the polar lignin cap in the exodermis is not affected; right panels = magnified image of vascular cylinder and endodermis layer. Small square right panel: Wild-type top view of CS and *slsgn3-1* top view of interrupted CS. White asterisks show the discontinuity of the CS in *slsgn3-1.* Brightness was adjusted in the magnified pictures, so the CS is more clearly visualized. Scale Bar = 50 with μm **E.** Translational fusions of the SlCASP2 and SlCASP1 proteins under the control of their respective promoters, to mCitrine, drive protein localization specifically in the tomato root endodermis and not in the exodermis. In longitudinal images, cell walls are stained by calcofluor (blue); lignin is stained by fuchsin (pink) and mCitrine is imaged in the GFP channel (green). Scale Bar = 50 μm **F.** Simultaneous mutation of *slcasp1* and *slcasp2* (*slcasp1/2*) showed no defect in endodermal CS or exodermal polar lignin cap deposition, relative to wild type. Lignin is stained by fuchsin (pink). Scale Bar = 50 μm. All CRISPR-generated mutant or reporter lines were generated by *A. tumefaciens* transformation unless otherwise noted.

Barrier function of the exodermal polar cap is dependent on lignin biosynthesis. Loss of function *atmyb36* mutants lack the CS and present ectopic lignification in endodermal cells (Kamiya et al. 2015; Reyt et al. 2021). Phylogenetic analysis revealed two possible tomato MYB36 homologs of Arabidopsis *AtMYB36* (At5g57620) -Solyc07g006750 and Solyc04g077260 (**Supplemental Figure 5A-B**). Solyc07g006750 is more phylogenetically related to *AtMYB36* (**Supplemental Figure 5B**) and its ribosome-associated transcript abundance is enriched in the endodermis (**Supplemental Figure 5C**) (Kajala et al. 2021). Solyc04g077260 is a recently reported homolog of *AtMYB36* (P. Li et al. 2018) (**Supplemental Figure 5A**). CRISPR-Cas9 hairy root mutants revealed that Solyc07g006750 (herein *SlMYB36*) but not Solyc04g077260 (herein *SlMYB36-b*) lacks a CS (**Supplemental Figure 5D**). *A. tumefaciens*-transformed CRISPR-Cas9 edited *slmyb36* mutant alleles lack a CS but have no changes in ground tissue radial patterning. In contrast to *atmyb36* mutants (Reyt et al. 2021; Kamiya et al. 2015), *slmyb36* mutants have minimal ectopic lignification within the endodermis. Neither *slmyb36* nor *slmyb36-b* mutants showed defects in the exodermal lignin cap (**Figure 2C**, **Supplemental Figure 5D**). These results are consistent with a conserved function for *SlMYB36* in endodermal differentiation, but not in exodermal differentiation.

AtCASP proteins act as a scaffold to guide lignin biosynthesis enzymes to the CS domain (Roppolo et al. 2011). We identified four putative tomato AtCASP homologs (Solyc02g088160, Solyc06g074230, Solyc09g010200, Solyc10g083250) (**Supplemental Figure 6**). Three of these have the extracellular loop1 (EL1) protein domain that is conserved in spermatophytes (Solyc06g074230, Solyc09g010200, and Solyc10g083250) (Roppolo et al. 2014) and were chosen for further study. Transcriptional GFP reporters demonstrated that *SlCASP1* (Solyc06g074230) is expressed primarily in the endodermis with some expression in inner cortical and exodermal cells, *SlCASP2* (Solyc10g083250) is primarily expressed in the endodermis and *SlCASP3* (Solyc09g010200) is expressed in the endodermis, cortex, and exodermis (**Supplemental Figure 4E**). SlCASP1 and SlCASP2 translational mCitrine reporters demonstrated that SlCASP1 and SlCASP2, but not the SlCASP3 proteins localize to the endodermal CS domain and not at the exodermal lignin cap domain in both hairy roots and A. tumefaciens transformed roots (**Figure 2E**, **Supplemental Figure 4E**). Hairy and *A. tumefaciens*-transformed roots with CRISPR-Cas9 edits in both SlCASP1 and SlCASP2 genes (*slcasp1slcasp2*) showed no phenotype in the endodermal CS or the exodermal polar lignin cap (**Figure 2F**), suggesting a high degree of functional redundancy among CASPs, similar to what is observed in *Arabidopsis* (Barbosa et al., 2022).

The AtSGN3 receptor kinase along with the AtCIF peptide controls the CS integrity surveillance system (Doblas et al. 2017; Nakayama et al. 2017). Phylogenetic analyses revealed one close homolog to *AtSGN3*/At4g20140, the Solyc05g007230 gene, and one to the *AtCIF1*/At2g16385 and *AtCIF2*/At4g34300 genes, Solyc01g109900 (also previously annotated as Solyc01g099895) (**Supplemental Figure 7 A-C**). Transcriptional GFP reporters for *SlSGN3* and *SlCIF1* showed the *SlSGN3* expression domain at the endodermis, inner cortex, and exodermis while the *SlCIF1* expression domain is in the vasculature as in *Arabidopsis* (Doblas et al. 2017; Nakayama et al. 2017) (**Supplemental Figure 4F**). Mutant alleles generated by *A. tumefaciens* and hairy root transformation via CRISPR-Cas9 editing of *SlSGN3* revealed an interrupted CS as observed for *Arabidopsis sgn3* (**Figure 2D; Supplemental Figure 4H**) (Pfister et al. 2014). The exodermal polar lignin cap is not affected in these mutants (**Figure 2D**; **Supplemental Figure 4H**). These data collectively demonstrate that *SlSHR*, *SlMYB36*, *SlCASP1*, *SlCASP2*, and *SlSGN3* are orthologs of *Arabidopsis* endodermal CS regulators, that endodermal differentiation is conserved between *Arabidopsis* and tomato and that tomato exodermal polar lignin cap differentiation is genetically distinct from endodermal differentiation.

### Characterizing two transcription factors that control exodermal differentiation

Mining of genes whose transcript abundance is enriched in the endodermis has been successful in identifying regulators of endodermal differentiation (Pfister et al. 2014; Y. Lee et al. 2013). We utilized this approach with the underlying hypothesis that transcription factors play a critical role in exodermal differentiation. Utilizing the exodermal enriched ribosome-associated transcript list (Kajala et al. 2021) we identified transcription factors from multiple families (MYB, NAC, AP2/ERF, MADS-BOX, HD-Zip, WRKY, HLH, Zinc finger C2H2, and HOMEOBOX) (**Supplementary Figure 8**). Of these, MYB transcription factors were particularly attractive candidates given their known role in endodermal barrier production, secondary cell wall biosynthesis, and lignin deposition (Shukla et al. 2021; Kamiya et al. 2015; Miyamoto, Tobimatsu, and Umezawa 2020; J. Liu, Osbourn, and Ma 2015). We selected eight MYB family transcription factors for CRISPR-Cas9 mutagenesis and compared lignin deposition in wild type (*R. rhizogenes*-transformed with no binary plasmid) relative to independent mutant alleles of each transcription factor (**Supplementary Table S1** and **Supplementary Figure 10**). We additionally mutated thirteen exodermal-enriched transcription factors (**Supplementary Table 1** and **Supplementary Figure 10**). A lignin phenotype was only observed for a mutant of a zinc finger C2H2 family (Solyc09g011120) transcription factor (five independent hairy root alleles) (Supplementary Table S1 and Supplementary Figure 11). Hairy root mutant alleles as well as *A. tumefaciens* transformed mutant alleles of Solyc09g011120 (herein *SlEXO1*) had ectopic lignin deposition in the form of a polar lignin cap in inner cortex cells in addition to a wild type exodermal polar cap (**Figure 3A**, **Supplementary Figure 11**). This phenotype suggests that *SlEXO1* represses polar lignin cap formation in inner cortical layers and thus restricts barrier formation to the exodermal cells. The root length of *slexo1-1* is additionally significantly shorter than wild type (Figure 3F).

**Figure 3.**
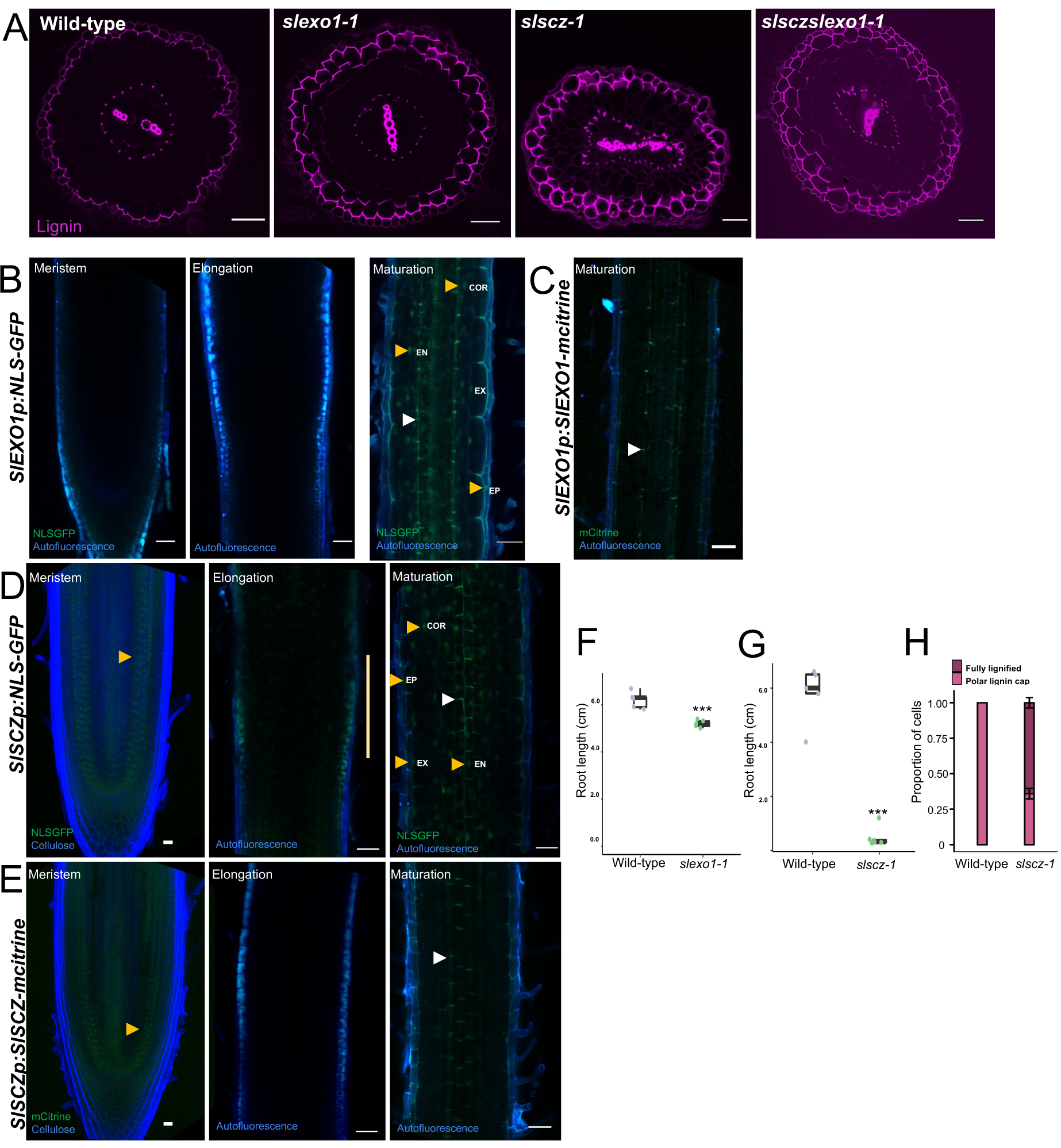
*SlEXO1* and *SlSCZ* repress lignification in the inner cortical layer(s). **A.** The *exo1-1* mutant allele has an additional lignin polar cap in the first inner cortical layers relative to wild type. The *slscz-1* mutant allele has perturbed lignification in the exodermis (either polar lignification, or non-polar lignification) and occasional lignification of the entire cell in the first inner cortical layer. The last exodermis/cortex layer (towards the vasculature) contains a polar lignin cap. Pink = fuchsin/lignin. The *slsczslexo1* double mutant has a similar phenotype as the single mutant *slscz-1*. Scale Bar = 50 μm **B.** *SlEXO1* transcriptional fusion (*SlEXO1*p::*SlEXO1-NLSGFP*). From left to right: Meristem, elongation, and maturation zones. *SlEXO1* is expressed in the epidermis, exodermis, inner cortex, and endodermis in the maturation zone. GFP (Green) and autofluorescence (Blue). Scale Bar = 50 μm. White and orange arrowheads point out the CS autofluorescence and NLS-GFP respectively. **C**. *SlEXO1* promoter fused to the SlEXO1 coding region and mCitrine to show the protein was not detected. White arrowhead points CS. **D.** *SlSCZ* transcriptional fusion (*SlSCZp*::SlSCZ-NLSGFP). From left to right: Meristem, elongation, and maturation. *SlSCZ* is expressed in the endodermis, cortex, and exodermis meristematic cells and the epidermis, exodermis, cortex, and endodermis in the maturation zone. Meristem images: Green= GFP. Blue = Calcofluor White. Elongation and maturation zone images: Green= GFP. Blue = autofluorescence. Scale Bar = 50 μm. White and orange arrowheads point to the CS autofluorescence and NLS-GFP respectively. **E.** *SlSCZ* promoter fused to the *SlSCZ* coding region and mCitrine demonstrates the protein is localized in the stem cell niche, the two inner cortical layers, and the exodermis. The SlSCZ protein was not detected at the elongation or maturation zone. Meristem images: Green= citrine. Blue = Calcofluor White. Elongation and maturation images: Green= citrine. Blue = autofluorescence. Images are captured from *R. rhizogenes* transformed roots. Scale Bar = 50 μm. White and orange arrowheads point to the CS autofluorescence and mCitrine respectively. **F.** Root length of wild type and the *slexo1* mutant (p-value=0.000465, one-way ANOVA). **G.** Root length of wild type and the *slscz-1* mutant (p-value=9.55e-07, ANOVA) (Signif. codes: 0 ‘***’ 0.001 ‘**’ 0.01 ‘*’ 0.05). **H**. Proportion of cells polarly and fully lignified in wild-type and *slscz-1*.

In addition to mining cell type resolution expression data, we selected candidate genes based on their function in root ground tissue patterning and specification in Arabidopsis. From these genes, a heat shock transcription factor *SCHIZORIZA* (*AtSCZ*/At1g46264) regulates asymmetric stem cell divisions and specification of root cortex cell identity (ten Hove et al. 2010; Pernas, Ryan, and Dolan 2010). In tomato, the cortex-endodermis initial cell gives rise to the endodermis, two inner cortical layers, and to the exodermis (an outer cortical layer) (Ron et al. 2013). As the exodermis is the outermost cortex cell layer in tomato, we hypothesized that a tomato homolog of *AtSCZ* could regulate exodermal differentiation. A single putative homolog of *AtSCZ* exists within the tomato genome, the Solyc04g078770 gene (herein *SlSCZ*) (**Supplemental Figure 9**). *A. tumefaciens*-generated CRISPR-Cas9 mutant alleles display non-polar lignification (lignin coating all faces of the exodermal cell) in most, but not all exodermal cells (**Figure 3H**). When lignin is non-polarized in the exodermal layer, lignin in the inner cortical layer is always polarly deposited (**Figure 3A**). Moreover, some independent mutant alleles of *slscz* in hairy roots have groups of lignified cells where lignification occurs across multiple consecutive inner cortical layers, again with the inner-most cell containing polar lignification (**Supplemental Figure 11B-D**). Sexually inherited mutations of *SlSCZ* (*A. tumefaciens*) are likely lethal as we were able to obtain only two independent mutant alleles (**Figure 3A**, **Supplemental Figure 11E**). *slscz-1* roots also have a short root phenotype, although much more extreme than *slexo1-1* as the root barely elongates after germination (**Figure 3G**). Ectopic lignin is also present in the endodermis and the vascular cylinder is asymmetrically organized (**Figure 3A**). Given the partial similarities between *slexo1-1* and *slscz-1*, we generated a double mutant to determine their genetic interaction, using the same guide RNAs to generate their respective mutations. Both hairy root and *A. tumefaciens*-transformed double mutant (homozygous) alleles showed a similar phenotype to the *slscz-1* mutant, with ectopic lignin in exodermis and inner cortex along with a shorter root (**Figure 3A**; **Supplemental Figure 11L**).

To understand the location of transcripts and proteins for these genes, we created transcriptional and translational reporter lines. NLS-GFP expression driven by the *SlEXO1* promoter was observed in the epidermis, exodermis, cortex, and endodermis from the maturation zone but not in the meristem or elongation zone (**Figure 3B**). The SLEXO1 protein was not detectable in hairy roots from a *SlEXO1* translational reporter *SlEXO1*::*SlEXO1*-mCitrine both with and without the 5’ and 3’ untranslated regions (Figure 3C). The *SlSCZ* transcriptional NLS-GFP reporter is expressed in the meristematic exodermis, cortex and endodermis and in the maturation zone in the epidermis, exodermis, cortex, and endodermis. We could not detect NLS-GFP in the elongation zone (**Figure 3D**). A translational reporter of *SlSCZ* fused to citrine showed that the SCZ protein is localized mainly at the meristematic exodermis, cortex and endodermis. The SCZ protein was not detected in the elongation or maturation zone (**Figure 3E**). The differences between the expression of *SlSCZ* and the localization of the SlSCZ protein suggest the role of post-transcriptional modification in determining the final location of SlSCZ activity (J.-Y. Lee et al. 2006).

### SlSCZ and SlEXO1 have partially overlapping downstream regulation

*SlEXO1* and *SlSCZ* both repress the lignification of inner cortical cell layers. Their double mutant phenotype further demonstrates that they genetically interact (**Figure 3A**). These data suggest that *SlEXO1* and *SlSCZ* may regulate common downstream regulatory pathways that control the synthesis of the exodermal lignin barrier. To address this hypothesis, we conducted transcriptome profiling of two independent alleles each of *slexo1* and *slscz* hairy root mutants. Principal component analysis revealed that the transcriptomes of each of these genotypes are transcriptionally distinct (**Figure 4A**). *Slscz* transcriptomes were more divergent from wild type than *slexo1* (**Figure 4A**), as also reflected by the higher number of differentially expressed genes (DEGs) that were identified for each mutant genotype relative to wild type (FDR test <0.01 Log2FC+/−0.6) with 263 genes representing common direct or indirect gene targets (**Figure 4B**). The majority of the *slscz*-DEGs were down-regulated relative to wild type (**Figure 4C**, **Supplementary Figure 12A-D**). From the common DEGs (**Figure 4B**, **Supplementary Figure 12D**) there is an enrichment of genes encoding peroxidases (Fisher’s exact test p-value = 5.92e-05), some receptor-like kinases and genes related to phenylpropanoid metabolism (**Figure 4D**). Gene Ontology categories associated with lignin and secondary cell wall biosynthesis were up-regulated in *slexo1*, consistent with the increased lignin in its inner cortical layer (**Figure 3A**, **Supplementary Figure 12B-E**). Conversely, genes associated with phenylpropanoid, and lignin biosynthesis were down-regulated in *slscz*, although genes associated with lignin catabolic processes, cell wall biogenesis, and cell organization were up-regulated (**Supplementary Figure 12C-E**). To confirm where these genes are expressed in wild type, we intersected these gene lists with those of tomato root cell types from Kajala et al., (2021). Genes differentially expressed in both the *slexo1* and *slscz* mutant alleles have enriched transcript abundance within wild type inner cortex, exodermis, and epidermal translatomes (Fisher exact test p-value < 0.05) (**Supplementary Figure 13A-B**) which support the likely repressive function of these transcription factors.

**Figure 4.**
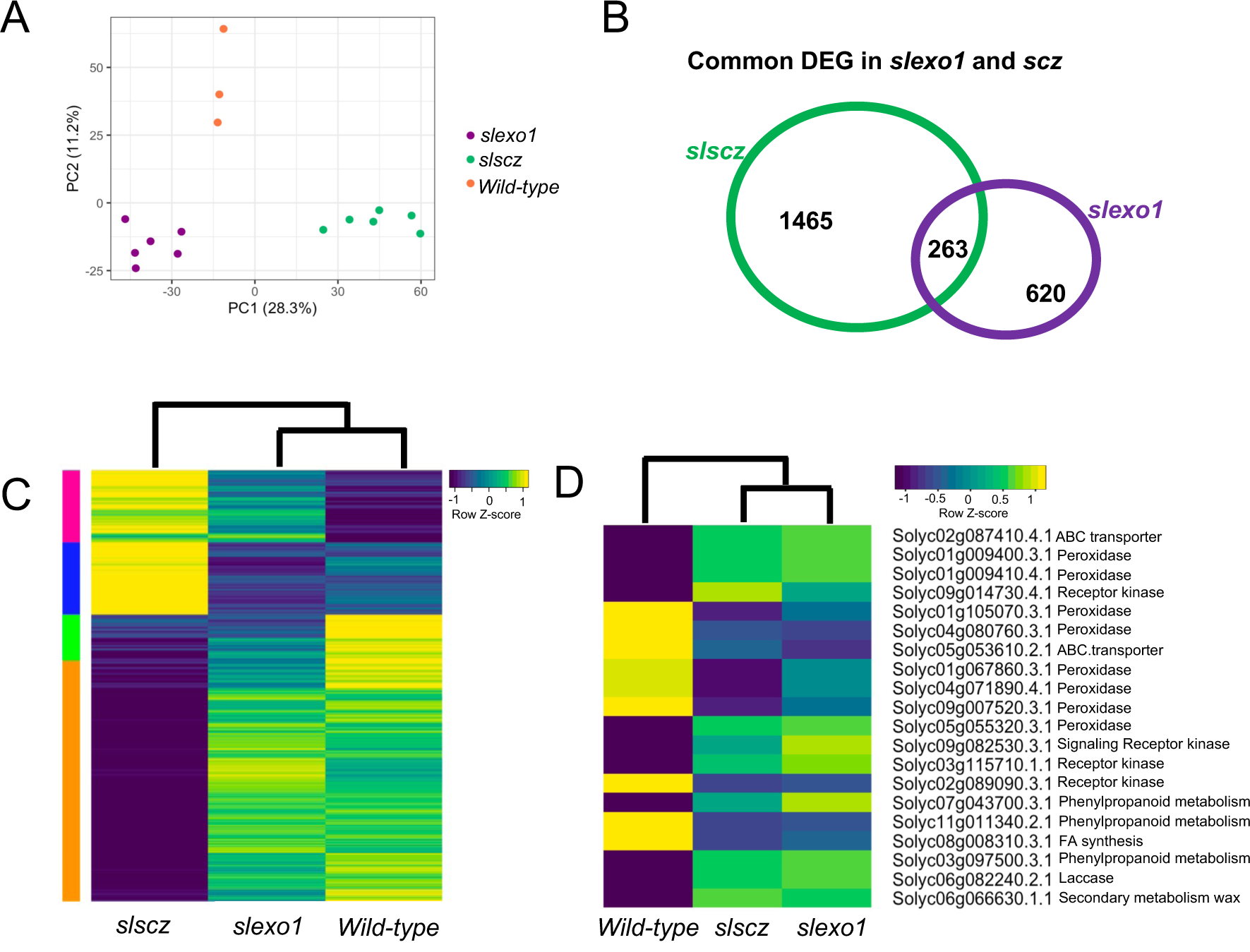
*SlEXO1* and *SlSCZ* transcriptionally regulate distinct and overlapping genes. **A.** Principal Component (PC) Analysis of the wild type (orange), *slscz* (green), and *slexo1* (purple) of *R. rhizogenes* transformed root transcriptomes. The first two dimensions contribute to 39.5% of the observed variation. The transcriptomes of these genotypes are distinct in PC1 and 2. **B.** Venn diagram indicating the common and uniquely differentially expressed genes in two independent *slexo1* mutant alleles and two independent *slscz* mutant alleles (FDR = 0.05; fold-change = 1.6) relative to wild type. **C.** Heatmap indicating significantly differentially expressed genes in each genotype (with *slexo1* and *slscz* representing differentially expressed genes in two independent alleles each) relative to wild type. Color intensity represents the row-normalized z-score. Rows are clustered with Pearson correlation and columns with Spearman correlation. **D**. Heatmap with the expression of some common genes from the Venn diagram from panel B. The selected genes are annotated as peroxidases, receptor kinases, ABC transporters and related to phenylpropanoid metabolism. There is a significant enrichment for peroxidases (Fisher exact test < 0.05).

### The exodermal and endodermal barriers are selective checkpoints for mineral ions

The endodermal Casparian strip plays a significant role in regulating the leaf ionome, consistent with the Casparian Strip as a selective barrier for mineral ion uptake to the shoot (Baxter et al. 2009; Hosmani et al. 2013; Pfister et al. 2014; Kamiya et al. 2015). The endodermis not only prevents mineral nutrients from entering the vasculature from the cortex but also prevents leakage back into the cortex of ions that are not translocated. In tomato, both the exodermis and endodermis have apoplastic barrier function leading to the question of whether each barrier has selective control of different ions. We tested this hypothesis by ionome profiling of the tomato shoot in the *slmyb36-1* mutant (CS absence but a wild-type exodermal lignin cap), the *exo1-1* mutant (no defects in the CS and a polar lignin cap in the first inner cortical layer), and the *slscz1-1* mutant (strong lignification in the exodermis and ectopic lignin in inner cortex cells and the endodermis) (**Figure 5A**). Leaves of four-week-old plants were analyzed for their elemental composition using inductively coupled plasma-mass spectrometry (ICP-MS) (**Figure 5**). Principal component analysis (PCA) revealed that *slmyb36-1* and *slscz-1* have different ionome profiles than wild-type, but the *exo1-1* mutant ionome is more similar to wild-type (**Figure 5B**). The levels of all ions tested are not significantly perturbed in *slexo1-1* relative to wild-type (two-way ANOVA, p<0.05). Leaves of *slmyb36-1* accumulate significantly increased sodium (Na), strontium (Sr), rubidium (Rb), calcium (Ca), magnesium (Mg), and lithium (Li) while molybdenum (Mo) is decreased in comparison with wild-type (**Figure 5C**; **Supplemental Figure 14**). The *slscz-1* ionome is the most perturbed relative to wild-type with increased accumulation of sodium (Na), rubidium (Rb), cobalt (Co), boron (B), and lithium (Li) and decreased levels of molybdenum (Mo), copper (Cu), manganese (Mn) and essential elements like potassium (K) and zinc (Zn) (**Figure 5C**; **Supplemental Figure 14**). Thus, the additional exodermal lignin barrier(s) in *slexo1-1*, *slscz-1*, and wild type does not completely compensate for the absence of the CS demonstrating that these barriers have unique and overlapping roles in selective mineral ion uptake.

**Figure 5.**
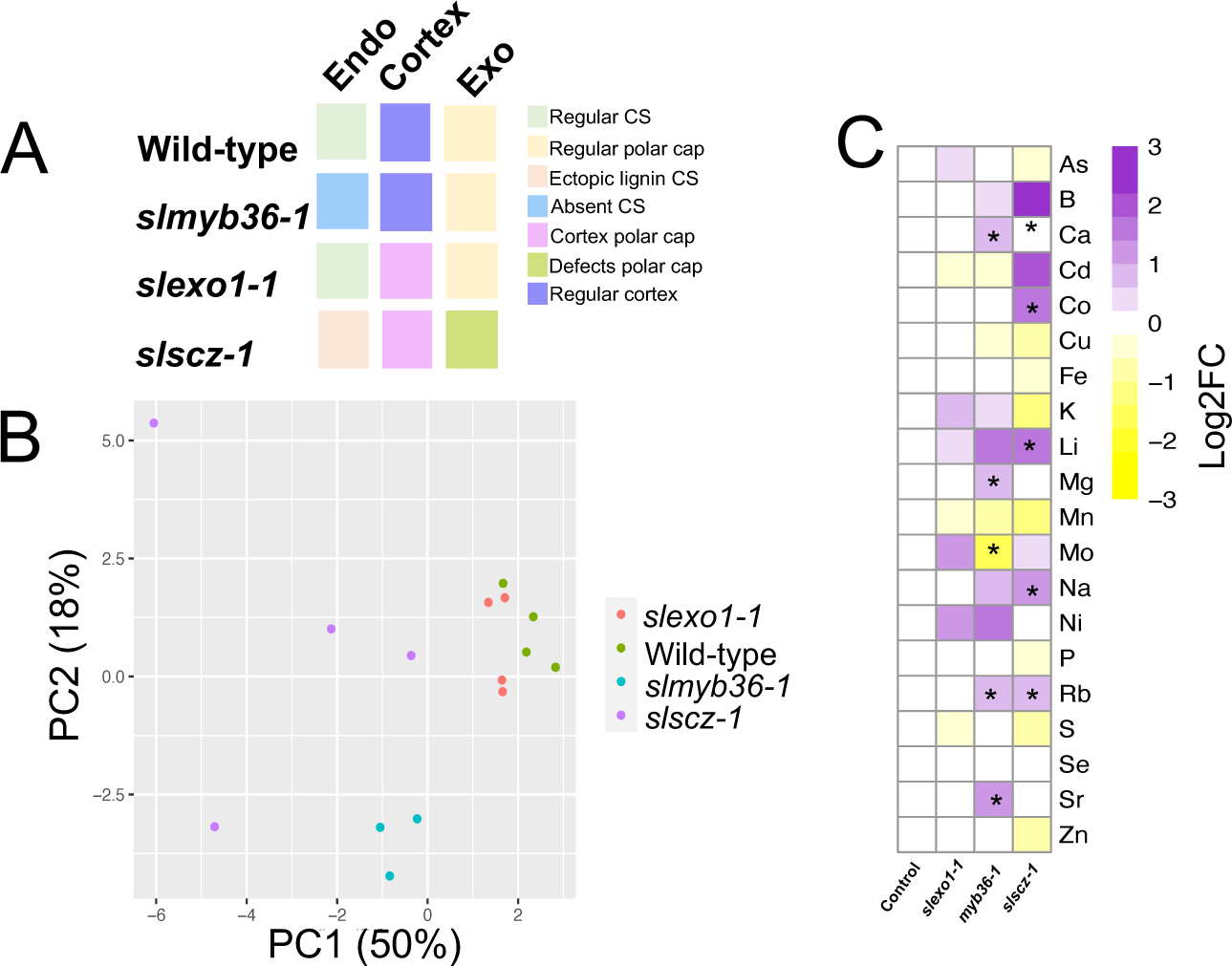
The exodermal lignin barrier does not compensate for the endodermal Casparian strip. **A.** Schematic representation of wild-type, *slmyb36-1*, *slexo1-1*, and *slscz-1* lignin barriers in the endodermis (Endo), cortex, and exodermis (Exo). **B.** Principal component analysis of 24 mineral ions within wild-type, *myb36-1, exo1-1*, and *slscz-1* mutant plants (n=4 for *slscz-1*, *slexo1-1*, and wild-type; n=3 for *slmyb36-1*). Considerable variation exists between the ionome of the *slscz-1* and *slmyb36-1* mutants. **C**. Log2 fold change of ions relative to wild type. Heatmap indicates the relative abundance of ions. *= statistical significance as determined by ANOVA with a post-hoc Tukey HSD test.

## DISCUSSION

The acquisition of multicellularity required the formation of intercellular barriers to facilitate communication and transport. The shape, composition, and location of these barriers inform their function. The best characterized of these barriers in plants is the endodermal CS, which is conserved across all angiosperms (Geldner 2013; Enstone, Peterson, and Ma 2002; C. Meyer and Peterson 2013; van Fleet 1961). As defined by Caspary and refined by others, this structure is formed of polymerized lignin that has impregnated the primary cell wall, as a precise and narrow belt in the center of the transversal and anticlinal walls (Alassimone, Naseer, and Geldner 2010; Enstone, Peterson, and Ma 2002; Naseer et al. 2012) (Caspary R. 1858; Caspary R. 1865). It fills up the entire space between two adjacent cells, including the middle lamellae, which means that the two cells are essentially fused with each other (Geldner 2013; Bonnett 1968; Haas and Carothers 1975; Karahara and Shibaoka 1992; Roppolo et al. 2011). The apoplastic barrier function of the CS is dependent upon its lignin composition (Naseer et al. 2012). In comparison, the root exodermis additionally forms apoplastic barriers that have been characterized for their functionality in many species (Peterson, Peterson, and Robards 1978; Peterson, Emanuel, and Wilson 1982; Peterson 1987; Lehmann et al. 2000). The shape and position of these barriers have also been elegantly described using histochemical stains and anatomical analyses, although many of these stains have not been able to distinguish between polymerized lignin and aliphatic suberin, nor do they facilitate ultrastructural observation (Enstone, Peterson, and Ma 2002) (Meyer and Peterson 2013). Here, we describe the developmental timeline by which a polymerized lignin cap seals the exodermal intercellular and epidermal-exodermal apoplastic space in a polar fashion. Although this structure impregnates the primary cell wall and serves as an apoplastic barrier dependent on its lignin composition, it is not centrally localized, as is the case for CS. How common is an exodermal barrier of this shape and composition across angiosperms? Although Perumalla et al. (1990) reported that several *Solanaceous* species possess an exodermis with a CS, our data clearly demonstrates otherwise. A similar exodermal polar structure composed of lignin and/or suberin was described in the literature for the seedless vascular plant *Sellaginella selaginoides* and for *Vinca minor*, the former of which is recalcitrant to cell wall digestion and thus fuses exodermal cells together (Damus et al. 1997; Perumalla, Peterson, and Enstone 1990). Barriers of other shapes have been described. The outer exodermis in *Iris germanica* has a reverse structure to the polar cap (called a Y-shape), while *Trillium grandiflorium*, sugarcane, onion, and barley roots have barriers sealing only the intercellular space of exodermal cells which resemble broadened CS (Lehmann et al. 2000; C. J. Meyer, Peterson, and Bernards 2011; Perumalla, Peterson, and Enstone 1990). Despite these differences in morphology, these structures appear to all act as apoplastic barriers, suggesting convergence.

The endodermis and exodermis have been compared in composition and apoplastic barrier function in tomato (Kajala et al. 2021; Cantó-Pastor et al. 2022). Two hypotheses can be formulated as to the mechanisms that give rise to the similarities in this first stage of exodermal and endodermal differentiation - that they are governed by similar developmental factors or that they are determined by distinct regulatory mechanisms. Analysis of tomato *SlSHR*, *SlMYB36*, *SlSGN3*, and *SlCASP* mutants demonstrates that these genes are conserved regulators of endodermis specification and differentiation, albeit with some differences. Mutation of tomato *slshr* reduces the number of ground tissue layers, and in part of the root, a CS is not evident. In the upper part of the root, however, a disorganized Casparian strip-like structure and ectopic lignin are observed in the mutant layer (the presumed second inner cortex layer). Furthermore, radial symmetry is perturbed in the ground tissue as well as the vascular cylinder. These additional phenotypes could reflect expanded function of *SlSHR* and differences in its regulatory network in other dicot species. The *slmyb36* mutant has a complete absence of lignin (**Figure 2C**), while the *atmyb36* mutant has ectopic lignin at cell corners which functions as a compensatory barrier. The phenotype observed in tomato could only be achieved in Arabidopsis when both *AtMYB36* and *AtSGN3* genes were mutated (Reyt et al. 2021). It is possible *SlMYB36* controls the expression of *SlSGN3*. A mutant phenotype was observed when the tomato *slsgn3* was mutated (**Figure 2D**), suggesting that a surveillance integrity mechanism does exist in tomato, although it is not necessary to compensate for a lack of endodermal CS. It is possible that this mechanism is not necessary for tomato as there is a cell type with an additional apoplastic barrier. Whether this remains true for other plant species with an exodermis remains to be seen. These data demonstrate that distinct regulatory mechanisms from the endodermis determine exodermal specification and deposition of the polar cap. The question of if other species with an exodermal barrier have co-opted the SHR/MYB36/SGN3 molecular regulators is still open.

Our extensive genetic screening for exodermal lignin cap regulators identified two transcription factors that repress lignin cap deposition in the inner cortex layer, with *SlSCZ* additionally regulating the polar deposition of lignin in the exodermis. The *AtSCZ* gene was previously identified as a regulator of cortex cell identity in Arabidopsis (ten Hove et al. 2010; Pernas, Ryan, and Dolan 2010). As there are multiple cortex cell layers (including the exodermis) in tomato, it is possible that *SlSCZ* has evolved to participate specifically during exodermis specification/differentiation, although defects in epidermal morphology are observed (**Figure 3A**). *SlSCZ* and *SlEXO1* reporter localization suggest that *SlSCZ* may regulate exodermal/inner cortical specification and differentiation, while *SlEXO1* regulates differentiation only. Collectively these data demonstrate active spatial restriction of exodermal differentiation to the outermost cortex cell layer. Plant species with a multiseriate (multiple-layered) exodermis (Peterson and Perumalla 1990) resemble these mutant phenotypes. Links with the environment and these barriers have been observed (C. J. Meyer, Seago, and Peterson 2009; C. J. Meyer, Peterson, and Steudle 2011; X. Li 2018) and it is intriguing to speculate that *SlSCZ* and *SlEXO1* enable this flexibility.

The relationship between these two factors can be inferred from the *slsczslexo1* double mutant. Here, the exodermal layer is largely fully lignified and when this occurs, the inner cortical layer has a polar lignin cap. The vascular cylinder is also elongated, and epidermal cells are misshapen. We propose two hypotheses regarding *SlSCZ* and *SlEXO1* genetic relationship. The double mutant phenotype is similar to that of *slscz* with respect to the complete lignification of the exodermal layer. One model is that the *slscz* phenotype is epistatic to that of *slexo1*, and *SlSCZ* is genetically downstream of *SlEXO1*. However, since *slscz* and *slexo1* phenotypes are not opposing, observing epistasis is not very conclusive. An equally plausible model is that these genes act additively, or in parallel pathways. Given the nature of these phenotypes, it is difficult to distinguish which model is the best fit. Despite these genes both being transcription factors, no changes in gene expression of the other factor were observed in their respective loss-of-function mutant alleles, although these data were not collected at cell type resolution. This suggests that the relationship between these factors is not via transcriptional regulation. Asymmetry in the vascular cylinder of both *slshr* and *slscz* was also observed, suggesting these two factors could coordinate aspects of vascular development. We utilized cell type-enriched gene lists to identify candidate genes that control exodermis development and to determine the domain of function of these genes, under the assumption their function is cell autonomous. These phenotypes in the vasculature (*slscz*, *slshr* and *slsczslexo1*), as well as the epidermis (*slscz*, *slsczslexo1*), raise the question of whether cell non-autonomous signals may further contribute to specification of the positioning of the polar cap. Additional clues to the factors that contribute to the positioning and polymerization of the cap were identified by transcriptional analyses of loss-of-function mutant alleles of both transcription factors. These include peroxidases and a laccase that could contribute to lignin polymerization as well as many signaling proteins including kinases and leucine-rich receptor kinases.

The tomato exodermal and endodermal barriers do act as a selective barrier for mineral ion uptake as previously observed (C. J. Meyer, Peterson, and Steudle 2011). No significant changes in element uptake were observed in the *slexo1-1* mutant, suggesting that although an internal lignin cap is observed in the inner cortical cell layer, it is not an additional functional barrier (**Figure 5B, C**). The ionomes of *slmyb36-1* and *slscz-1* were quite different from wild type and each other (**Figure 5B**) demonstrating extensive differences in ion uptake in a genotype with an absent CS relative to that of a genotype with multiple lignified cortical layers. An increase in Cadmium (Cd), Cobalt (Co), and Lithium (Li) were solely observed in *slscz-1*, suggesting that specific transporters may be enriched in cortical cells with this barrier type (**Figure 5C**). Calcium (Ca), magnesium (Mg), and strontium (Sr) were enriched in the ionomes of *slmyb36*, demonstrating that the CS likely restricts their passage to the vasculature (**Figure 5C**) and the lack of compensation by either *slexo1-1* or *slscz-1* indicates their inability to compensate for this missing barrier. This contrasts with observations for silicon and manganese uptake in rice where their transporters are expressed in both the exodermis and the endodermis (Ma et al. 2007) (Sasaki et al. 2012). Both sodium (Na) and rubidium (Rb) over accumulate in *slmyb36-1* and *slscz-1*. Perhaps this could be explained by the CS restricting their uptake, and the extra barriers in *slscz-1* promoting their uptake. The function of these barriers may additionally have an influence in response to the abiotic and biotic environment. In the *slmyb36-1* mutant, there are no changes in essential elements as was observed in *atmyb36* (Reyt et al. 2021). A mutant with a complete absence of the exodermal barrier remains to be identified and is necessary to completely delineate the roles of each of these barriers. A breeding advantage may be conferred by the consecutive lignin caps of *slexo1-1* (exodermis and inner cortex) in toleration of higher ion concentrations in the soil matrix or rhizosphere as results from a restriction in water availability or high salinity as was reported for a species with a multilayered exodermis (C. J. Meyer, Peterson, and Steudle 2011). In addition, the presence or absence of the exodermal lignin cap and the endodermal CS may influence the uptake of specific microbes or other biotic organisms which could be selectively modulated for increased pathogen resistance (Kawa and Brady 2022). We also hypothesize that the differences between the two exodermis mutants ion phenotypes could be explained not just by the presence or absence of ectopic lignin in exodermis and inner cortex, but also by the amount and composition of lignin in these cell types, as was proposed for corner lignification in the *atmyb36-1* mutant (Reyt et al. 2021). Also, the *slscz* mutant has additional phenotypes like a shorter root that can also influence these distinct ionome profiles. Finally, these distinct barriers may provide mechanical support to the root system as it grows through the soil, as described for lignin in other plant tissues (Polo et al. 2020; Q. Liu, Luo, and Zheng 2018; Zhao et al. 2020; Emonet and Hay 2022).

To conclude, we have described a polarly localized lignin barrier in the tomato root exodermis which functions as an apoplastic barrier and controls selective mineral ion uptake. We have elucidated a gene set that controls its developmental regulation, and which is distinct from endodermal regulation. Transcriptomic analyses of these mutants identified several factors that may play a role in the positioning and polymerization of the exodermal lignin cap.

## MATERIAL AND METHODS

### Plant Material and Growth Conditions

*S. lycopersicum* (cv. M82, LA3475) seeds were surface sterilized with 70% ethanol for 5 minutes and then treated with commercial bleach (70%) for 20 minutes. Seeds were subsequently washed three times with deionized water. Seeds were transferred to 12 cm x 12 cm plates containing 4.3g/L Murashige and Skoog medium (Caisson), 0.5g/l MES pH 5.8 and 10 g/L agar (Difco). Plates were placed in a 23°C growth chamber for seven to ten days with 16 hours of light and 8 hours of dark per day. All sections are from 5-6 days-old plants.

### Molecular Cloning

For the transcriptional fusions, a region of 2-3Kb upstream of the ATG was selected as the promoter region. This PCR fragment was subcloned into D-TOPO entry vector (Fisher scientific Cat no 450218). The D-TOPO clone containing the promoter fragment was recombined using an LR reaction into the vector pMK7FNFm14GW (Karimi et al. 2007) using the LR gateway cloning system (Thermo Fisher).

For the translational fusions, a region of 2-3Kb upstream of the ATG was selected as the promoter region. This fragment was cloned in 5 primeTOPO (Thermo Fisher scientific Cat no 591-20), the CDS was cloned in D-TOPO (Fisher scientific Cat no 450218) and the mCitrine cloned in P2P3 gateway vector (The P2P3 with mCitrine vector was kindly provided by the lab of Niko Geldner). All three vectors were recombined using multisite LR Gateway into the destination vector pB7m34GW (Karimi, De Meyer, and Hilson 2005).

### Rhizobium rhizogenes *Transformation*

“Hairy root” transformation with *R. rhizogenes* (ATCC 15834) was conducted according to (Ron et al. 2014). Cotyledons from seven to ten-day-old plants were cut and immersed immediately in a suspension of *R. rhizogenes* containing the desired binary vector. Cotyledons were then floated on sterile Whatman filter paper and co-cultivated in the dark on MS agar plates (1X vitamins, 3% sucrose, 1% agar) for three days at 25°C in the dark, without antibiotic selection. Cotyledons were then transferred to a selective plate (MS with cefotaxime; 200 mg/L; 50 mg/L kanamycin). 30 independent antibiotic-resistant roots were subcloned for each transgenic line for future analyses with Cefotaxime + Kanamycin for 2 rounds of subcloning and maintained in media with Cefotaxime after that period.

### Agrobacterium tumefaciens *Transformation*

The UC Davis Plant Transformation Facility generated transgenics using *A. tumefaciens* and tissue-culture-based protocols.

### Histochemistry and Imaging

Hairy roots of transcriptional fusions (*SlCASPs, SlSGN3, SlCIF, SlEXO1*, and *SlSCZ*) were imaged using a Zeiss LSM700 confocal microscope (water immersion, 20X objective) with excitation at 488 nm and emission at 493–550 nm for GFP for mCitrine and excitation at 555 nm and emission at 560–800 nm for autofluorescence. For the SlSCZ:GFP transcriptional fusion and the *SlSCZ*-mCitrine translational fusion images within the meristematic zone, hairy roots were cleared in Clearsee solution for 2 weeks (Ursache et al. 2018), stained with Calcofluor White for 30 minutes, washed twice and visualized using a Zeiss Observer Z1 fluorescent microscope. The same protocol was used to image the stable lines of the translational fusion constructs (*SlCASP1p*::SlCASP1-mCitrine and *SlCASP2p*::SlCASP2-mCitrine) and the *slcasp1slcasp2* stable CRISPR mutant.

For root sections and their staining, 1 cm root segments were embedded in 3% agarose and sectioned using a vibratome (Leica VT1000 S) to produce 150-200 μM thickness sections. The sections were then cleared and stained in Clearsee solution with Basic Fuchsin (Fisher scientific Cat no 632-99-5) for 30 minutes followed by two washing steps in Clearsee only. As indicated in the figures, in some cases, the Basic Fuchsin staining was followed by Calcofluor White (PhytoTechnology laboratories Cat no 4404-43-7) staining for 30 minutes and followed by two washing steps in Clearsee solution. Imaging of cellulose and lignin was performed using a Zeiss LSM700 Laser scanning microscope with the 20X objective. For Basic Fuchsin: 550-561 nm excitation and 570-650 nm detection were used. Root samples were mounted in ClearSee (Ursache et al., 2018) and scanned and imaged using the Zeiss Observer Z1 (20X or 40x objective). The following settings were used: Basic Fuchsin: 550-561 nm excitation and 570-650 nm detection for Basic Fuchsin and 405 nm excitation and 425-475 nm detection for Calcofluor White.

### ICP-MS

Seeds from *myb36-1, slexo1-1, slscz-1*, and wild-type M82 were germinated in 1% MS + 1 % sucrose square plates, and then 6 days after germination they were transferred to pots in soil. Four plants per genotype were randomized on a tray and watered two times a week with fertilized water for one month. The same portion of the compound leaf was collected from each plant and dried in a falcon tube in a 60C oven for 12 hours. Dried tissue was homogenized with a mortar and pestle and the dried powder was weighed for further analyses. The powder was digested in concentrated nitric acid for 3 hours at 100C. After digestion, nitric acid was evaporated at 80-100C. The same volume of 2% Nitric acid was added to each sample. The standards for the 20 elements were from Sigma Aldrich (01969-100ML-F, 01932-100ML;, 19051-100ML-F, 36379-100ML-F, 30329-100ML-F, 68921-100ML-F, 06335-100ML, 12292-100ML, 30083-100ML-F, 74128-100ML, 68780-100ML, 00462-100ML;, 28944-100ML-F, 38338-100ML, 01444-100ML, 18021-100ML, 50002-100ML, 75267-100ML) and Fisher scientific (CLZN22M, CLFE2-2Y) and were prepared in a dilution series. Samples were analyzed in a Perkin Elmer ICP-MS. Calibration curves were made using the standards and the amount (part per million) was calculated based on the calibration curve. Data analyses were performed in R for the PCA plot (prcomp package) and the heatmap (pheatmap from Biostrings package). A two-way ANOVA statistical test was performed (p-value<0.05) to identify elements significantly different from wild-type. The relative amount for each element per genotype in the Heatmap was calculated as the log2 fold change from the average of 3-4 replicates.

### Apoplastic Tracer Assays

Seeds of M82 were grown in 1% MS + 1% sucrose. Four days after germination, plants were transferred to new MS square plates with/without 200 μM of piperonylic acid (Sigma P49805-5G) and 200 μM piperonylic acid plus 10mM monolignols (Sigma Aldrich 223735-100MG,404586-100MG) for 24 hours in the dark, with DMSO as the solvent. Three cm segments from the root tip including the root meristem were exposed to 15 μg/mL PI (P4170; Sigma) in the dark at 30°C for 1 h. The roots were washed in water and one cm segments from the root tip were embedded in 3% agarose and made sections of 150-200 μM using a vibratome (Leica VT1000 S). Confocal Laser Scanning microscopy was performed on a Zeiss LSM700 confocal with the 20X objective, at 405nm excitation and 600-650 nm detection.

### Transmission Electron Microscopy

Tomato roots were fixed in 2.5% glutaraldehyde solution (EMS, Hatfield, PA) in phosphate buffer (PB 0.1 M [pH 7.4]) for 1 hour at room temperature and subsequently fixed in a fresh mixture of osmium tetroxide 1% (EMS) with 1.5% potassium ferrocyanide (Sigma, St. Louis, MO) in PB buffer for 1 hour at room temperature. The samples were then washed twice in distilled water and dehydrated in acetone solution (Sigma, St Louis, MO, US) in a concentration gradient (30% for 40 minutes; 50% for 40 minutes; 70% for 40 min and 100% for 1 hour 3 times. This was followed by infiltration in LR White resin (EMS, Hatfield, PA, US) in a concentration gradient (33% LR White 33% in acetone for 6 hours; 66% LR White in acetone for 6 hours; 100% LR White for 12 hours two times) and finally polymerized for 48 hours at 60°C in an oven in atmospheric nitrogen. Ultrathin sections (50 nm) were cut transversely at 2, 5, and 8 mm from the root tip, the middle of the root, and 1 mm below the hypocotyl-root junction, using a Leica Ultracut UC7 (Leica Mikrosysteme GmbH, Vienna, Austria), picked up on a copper slot grid 2×1mm (EMS, Hatfield, PA, US) and coated with a polystyrene film (Sigma, St Louis, MO, US).

Visualization of lignin deposition in the exodermis and the CS was performed using permanganate potassium (KMnO_4_) staining (Hepler et al., 1970). The sections were post-stained using 1% KMnO_4_ in H_2_O (Sigma, St Louis, MO, US) for 45 minutes and rinsed several times with H_2_O.

Micrographs and panoramic images were taken with a transmission electron microscope FEI CM100 (FEI, Eindhoven, The Netherlands) at an acceleration voltage of 80kV with a TVIPS TemCamF416 digital camera (TVIPS GmbH, Gauting, Germany) using the software EM-MENU 4.0 (TVIPS GmbH, Gauting, Germany). Panoramic images were aligned with the software IMOD (Kremer et al, 1996)(Kremer, Mastronarde, and McIntosh 1996).

### Transcriptome Profiling and Data Analysis

For the RNA-seq analyses three cm of the same developmental stage of hairy roots from *slscz-hr5*, *slscz-hr12*, *slexo1-hr6*, *slexo1-hr7* and wild-type M82 transformed with R.rhizogenes with no vector in three biological replicates. RNA was extracted with the Direct-zol RNA MiniPrep Plus kit (Neta Scientific Cat no RPI-ZR2053). cDNA libraries were made with the QuantSeq 3′ mRNA-Seq Library Prep kit from Lexogen (Cat no 015 QuantSeq FWD 3’ mRNA-Seq Library Prep Kit – with single indexing). Sequences were pooled, trimmed, and filtered using Trimmomatic (Bolger, Lohse, and Usadel 2014). Trimmed reads were pseudo-aligned to the ITAG4.1 transcriptome (cDNA) (Tomato Genome Consortium, 2012) using Kallisto (v0.43.1)(Bray et al. 2016), with the parameters -b 100–single −l 200 −s 30, to obtain count estimates and transcript per million (TPM) values. Differentially expressed genes (DEGs) were detected with the limma R package, using normalized CPM values as required by the package (Ritchie et al. 2015). CPM values were normalized with the voom function (Law et al., 2014) using TMM normalization. The functions lmfit, contrasts.fit, and ebayes were used to fit a linear model and calculate differential gene expression between the different contrasts. Genes with a log2 fold change (FC) value +/- 0.6 and FDR adjusted P value (adj.P.Val) ≤ 0.01 were considered as differetially expressed. The fdr method was used to control the false discovery rate (FDR) (Benjamini and Hochberg 1995).

### Generation of CRISPR-Cas9 edited constructs: hairy root mutant screen

Target guide RNAs were designed using the CRISPR-PLANT web tool (https://www.genome.arizona.edu/crispr/CRISPRsearch.html). In cases where CRISPR-PLANT did not specify at least three guides with GC content between 40 to 60%, guides were designed with CRISPR-P V2 (http://crispr.hzau.edu.cn/cgi-bin/CRISPR2/CRISPR), using the U6 snoRNA promoter with < 3 mismatches within the target gene coding sequence. Genomic sequences (ITAG3.2) were retrieved from Phytozome (https://phytozome-next.jgi.doe.gov/) and gene maps were constructed with SnapGene. Primers for genotyping were designed with Primer-BLAST software (https://www.ncbi.nlm.nih.gov/tools/primer-blast/). Primer specificity was checked against the *Solanum lycopersicum* using blast in Phytozome. The guide RNA was cloned in a method adapted from Lowder et al. (2015). In summary, oligos containing the sgRNA PAM sequence were phosphorylated and ligated into pYPQ131-3 vectors and recombined into p278 via Gateway cloning. A p278 vector containing all 3 gRNA expression cassettes was then recombined by Gateway into a pMR286 vector containing Cas9 and Kan resistance expression cassettes (Bari et al. 2019). The final CRISPR vector was introduced into *Rhizobium rhizogenes* (hairy roots) and *Agrobaterium tumefaciens* (stable lines) to generate transgenics.

### Phylogenetic tree construction

The following pipeline from Kajala et al., 2021 was utilized. Forty-two representative proteomes were downloaded from Phytozome, Ensembl, or consortia sites depending on availability. These include early diverging taxa, and broadly representative taxa from angiosperms. Next, blastp (Madden, 2013) was used to identify homologous sequences within each proteome based on a sequence of interest, with options “-max_target_seqs 15- evalue 10E-6-qcov_hsp_perc 0.5 - outfmt 6.” To refine this set of sequences, a multiple sequence alignment was generated with MAFFT v7 (Katoh and Standley, 2013) (option–auto), trimmed with trimal (Capella-Gutiérrez et al., 2009) with setting “-gappyout,” and a draft tree was generated with FastTree (Price et al., 2010). A monophyletic subtree containing the relevant sequences of interest was selected and more distantly related sequences were removed from the list of sequences. Tree construction methodology was informed by Rokas, 2011. For the final trees, MAFFT v7 using L-INS-i strategy was used to generate a multiple sequence alignment. Next, trimal was used with the -gappyout option. To generate a phylogenetic tree using maximum likelihood, RAxML was used with the option −m PROTGAMMAAUTO and 100 bootstraps. Finally, bipartitions with bootstrap values less than 25% were collapsed using TreeCollapserCL4 (http://emmahodcroft.com/TreeCollapseCL.html). The resulting trees were rooted on sequences from the earliest-diverging species represented in the tree.

## DATA AND RESOURCE AVAILABILITY

CRISPR-generated mutant lines are available upon request. Timing is dependent upon obtaining phytosanitary certificates according to seed import regulations of the country of destination and associated costs.

## SUPPLEMENTAL FIGURES

**Supplemental Figure 1.**
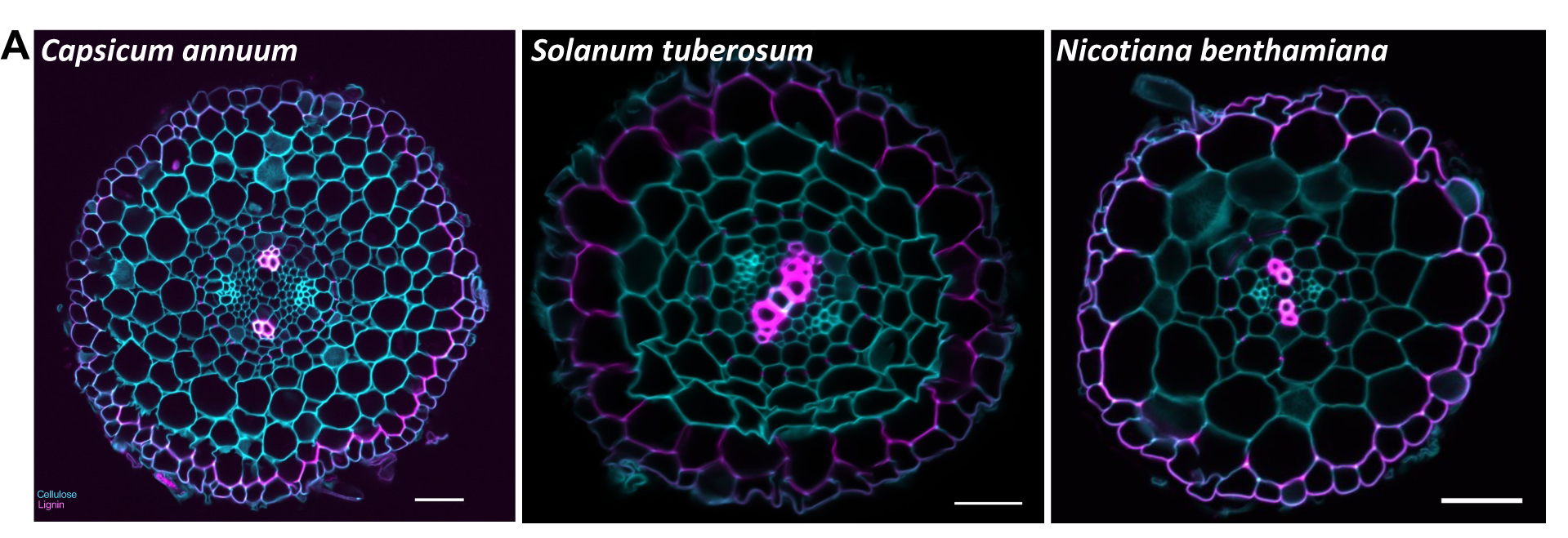
The polar lignin cap is conserved in the Solanaceae family. **A**. Cross sections of Solanum species and *Nicotiana benthamiana* stained with fuchsin and calcofluor. Left panel: *Capsicum annuum* variety Alliance bell pepper. Middle panel: *Solanum tuberosum* group Andigenum. Right panel: *Nicotiana benthamiana*.

**Supplemental Figure 2.**
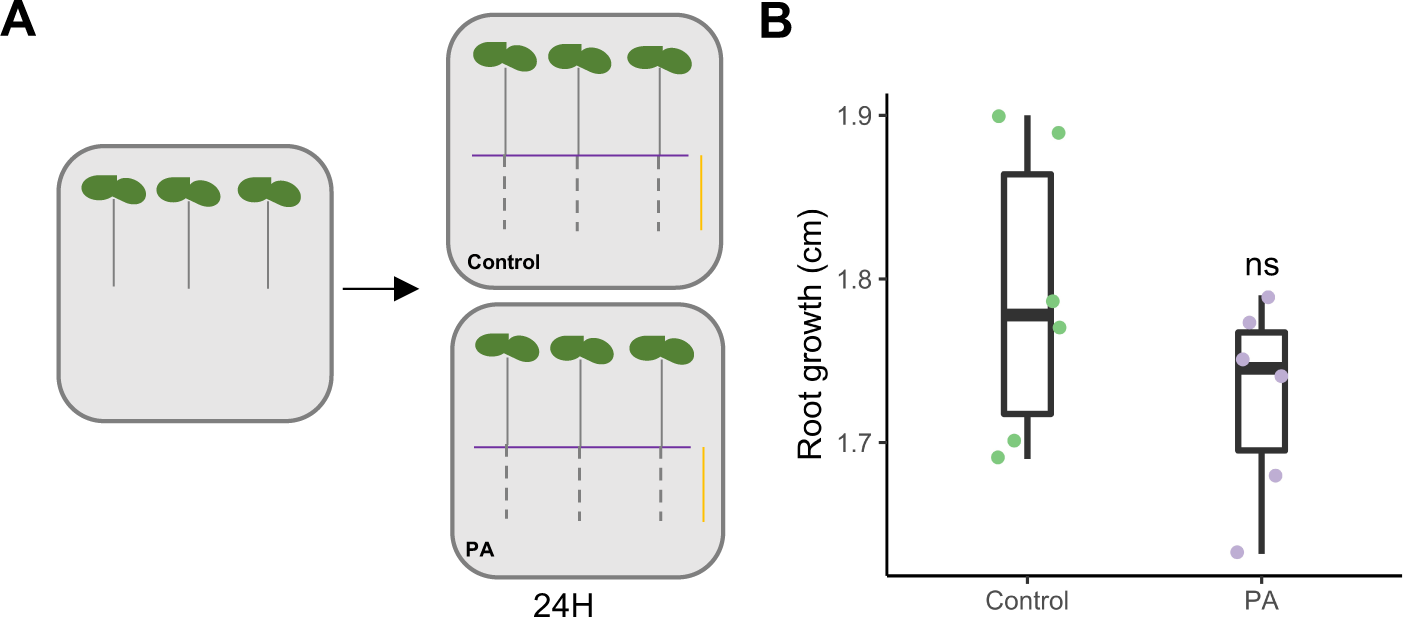
The piperonylic acid concentration tested for lignin biosynthesis inhibition does not affect root growth. **A.** Schematic representation of the experimental design to test the PA. 4-day-old plants after germination were transferred to MS media + 1% sucrose with and without 200 μM of PA. After 24 hours, root growth was measured for the part of the root that grew after transferring. Orange horizontal bar = newly grown root after transferring. **B**. Root growth after 24H in control and PA treatment for the new root zone after transferring (statistical significance test determined by ANOVA p-value 0,05).

**Supplemental Figure 3.**
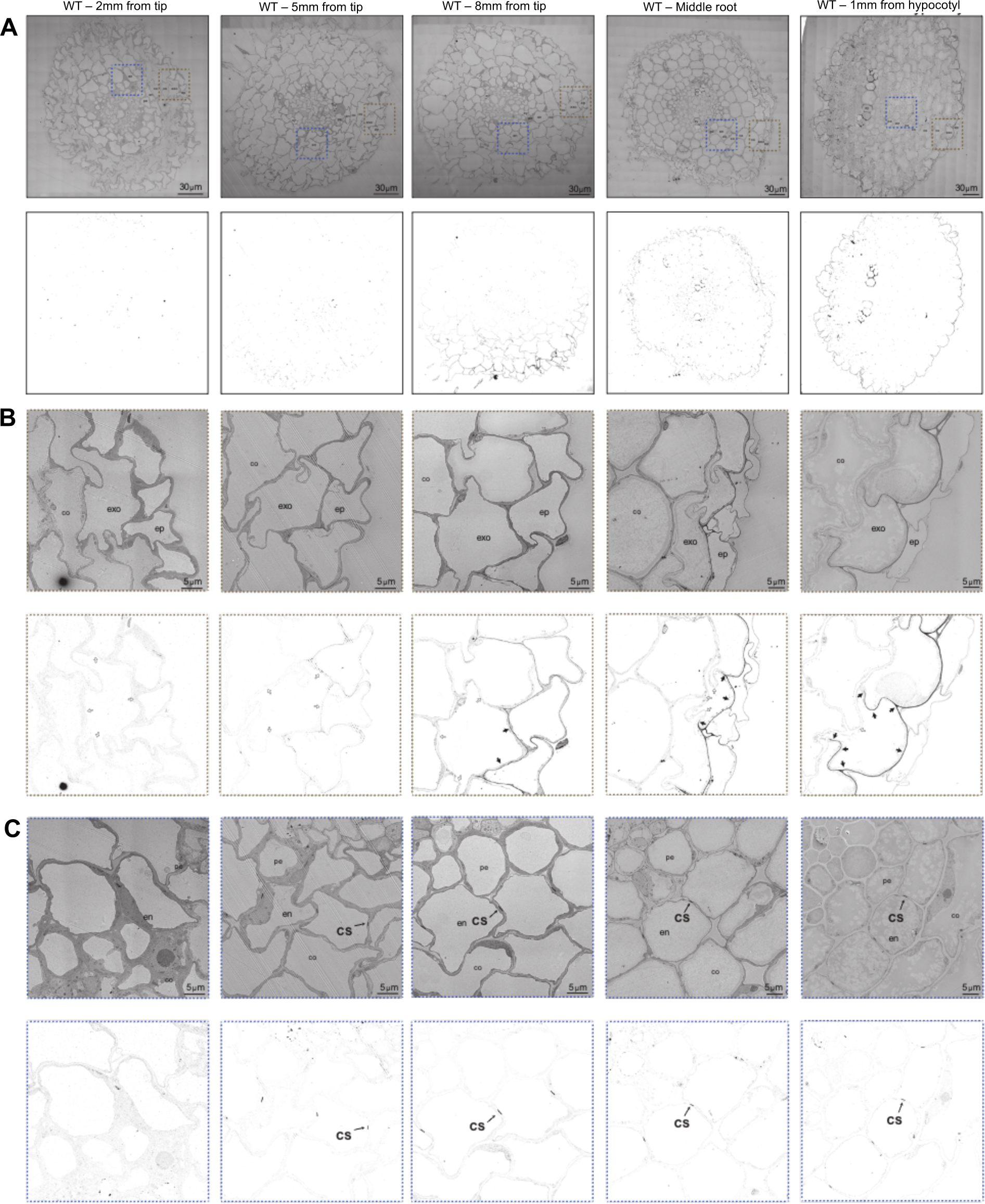
Ultrastructural lignin deposition in the exodermis and endodermis. **A.** TEM panoramas from wild-type root cross-sections at 2, 5, and 8 mm from the tip, a middle section of the root, and at 1mm from hypocotyl stained with KMn04. Dark deposits indicate electron dense Mn02 precipitation caused by reaction with lignin. Lower panels show the same pictures with an adapted contrast that highlights the dark deposits. Co = cortex, exo = exodermis, ep = epidermis, en = endodermis, xy = xylem, scale bars=30 μm. B. Close-up of the exodermis area from panels in A (zone defined with brown dashed lines). Lower panels show the same pictures with an adapted contrast that highlights the dark deposits. Dark arrows highlight lignin deposition in the exodermis cell wall. White arrows highlight the absence of lignin in the exodermis cell wall. Lignin deposition begins at 8mm from the root tip and is oriented toward the epidermis. Co = cortex, exo = exodermis, ep = epidermis, scale bars=5 μm. C. Close-up of the endodermis area from panels in A (zone defined with blue dashed lines). Lower panels show the same pictures with an adapted contrast that highlights the dark deposits. The CS is fully lignified at 5mm from the root tip. co = cortex, en = endodermis, pe = pericycle, cs = Casparian strip, scale bars=5 μm.

**Supplemental Figure 4.**
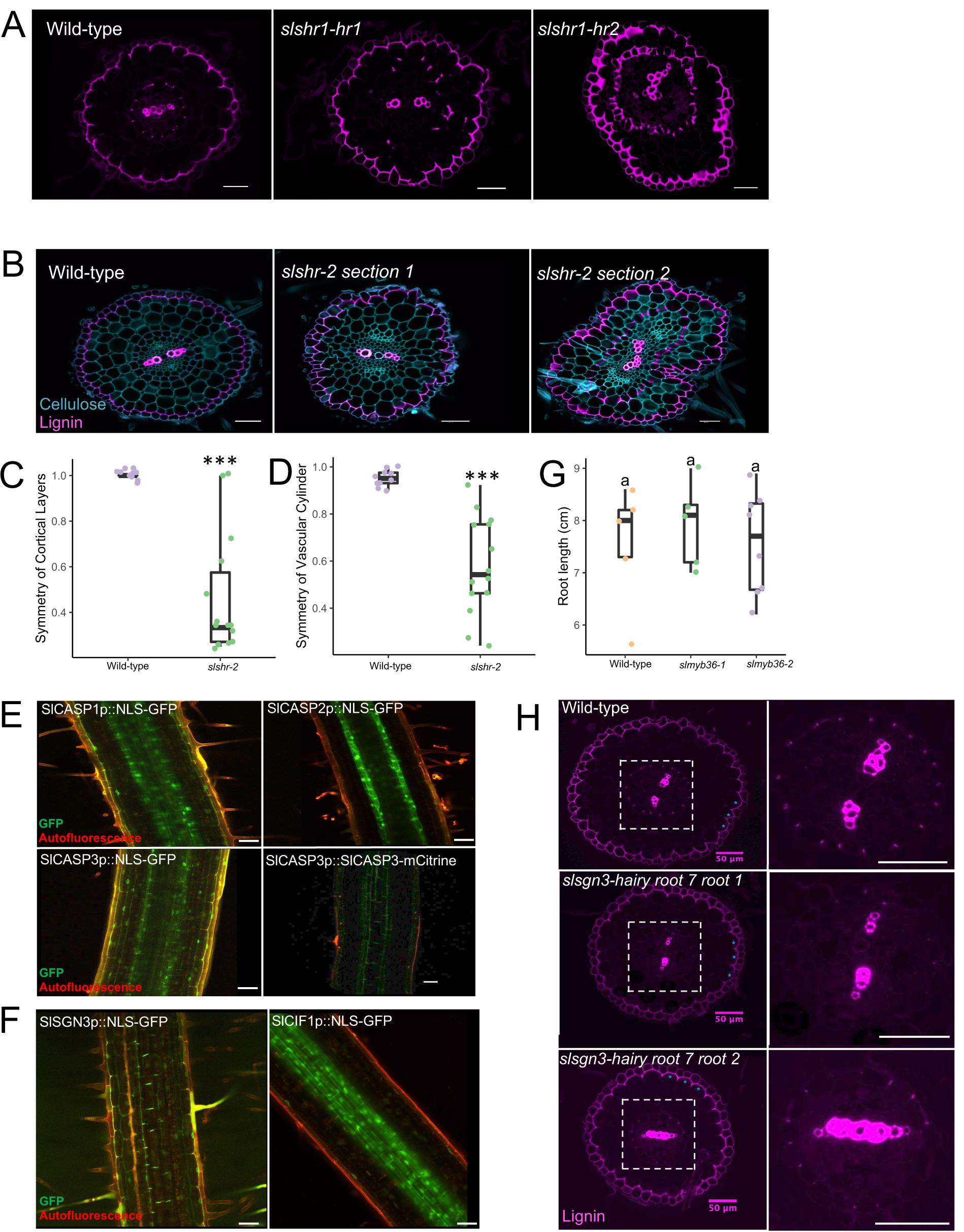
Endodermal regulators do not regulate exodermal differentiation. **A.** Hairy root cross-sections stained with fuchsin of wild-type (transformed with *R.rhizogenes* with no binary plasmid) and *shr-1* mutants from 2 independent hairy root lines. Left panel = wild-type, middle panel = *slshr1 hairy root Line 1 (slshr1-hr1*), right panel = *slshr1 hairy root line 2 (shr1-hr2*). **B.** Cross sections of wild-type and the *shr-2* mutant allele (*A. tumefaciens*-transformed) from two different parts of the root - 8 mm from the tip (middle) and 10 mm from the tip (right). The mutant layer undergoes ectopic lignin deposition in more mature regions of the root. **C.** Left panel: Radial symmetry of the cortical layers (including the exodermis). This is calculated as the minimum number of cortex layers observed divided by the maximum number of cortex layers in wild type and *shr-2* mutant roots (p-value=2.86e06, *=statistical significance as determined by ANOVA n=10-14. **D**. Symmetry of the vascular cylinder was calculated as the minimum distance across the center of the vascular cylinder divided by the maximum distance across the center of the vascular cylinder (p-value=1.36e05, *=statistical significance as determined by ANOVA n=10-14). **E.** Transcriptional fusions for *CASP1-3* genes to GFP and a translational fusion of *SlCASP3* to mCitrine. **F.** *SlSGN3* and *SlCIF2* transcriptional fusion to GFP. **G**. Root length in cm of two *slmyb36* independent alleles. Statistical significance as determined by ANOVA with a post-hoc Tukey HSD test. n=5 **H.** Mutation of *SlSGN3* in the hairy root line 7 results in an absent CS, or in a perturbed CS in the radial axis. Left panels = whole root section stained with fuchsin. Blue asterisks show the polar lignin cap in the exodermis is not affected.; right panels = magnified image of vascular cylinder and endodermis layer. Scale Bar = 50 μm.

**Supplemental Figure 5.**
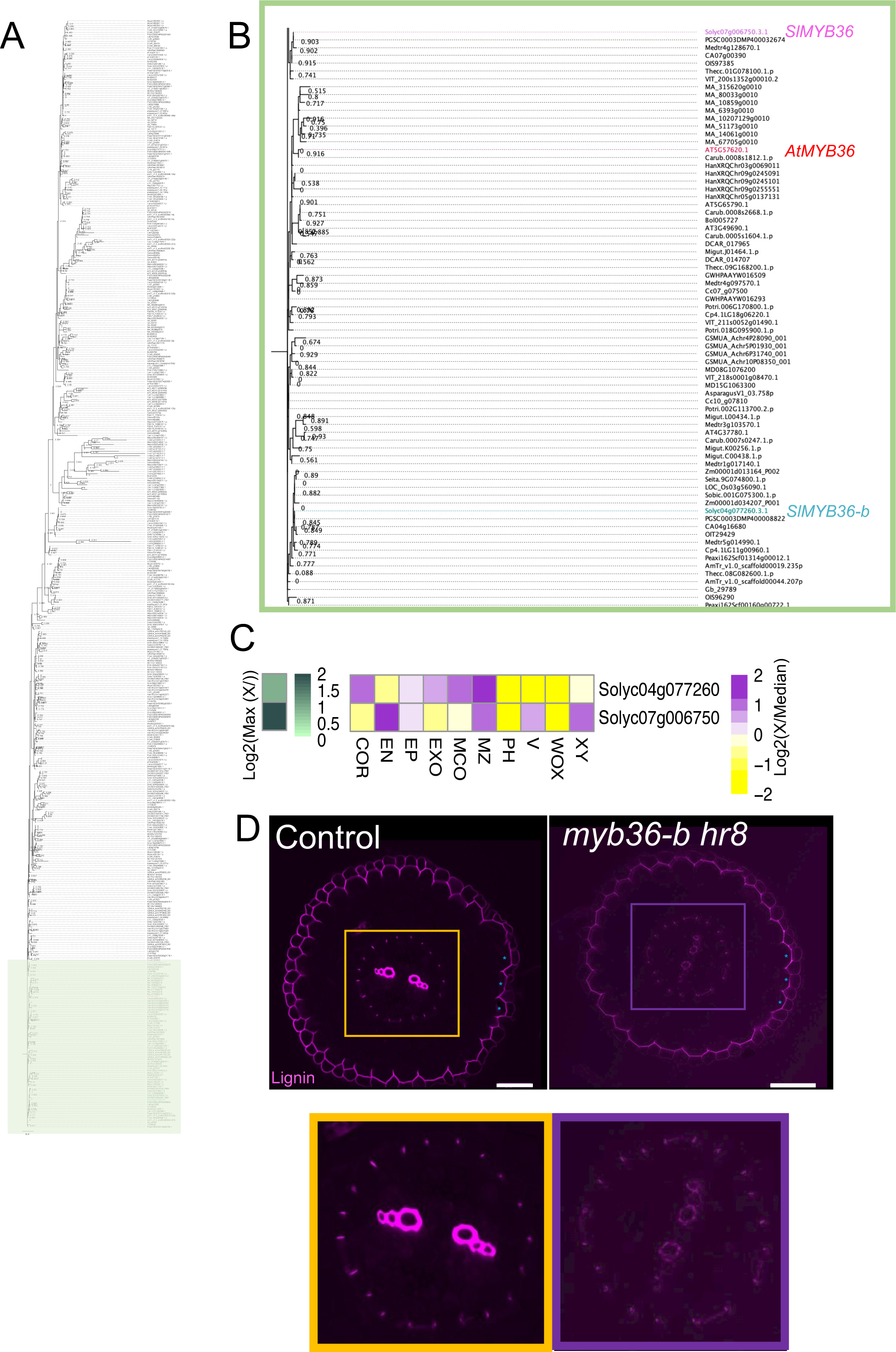
A previously identified *SlMYB36-*like gene is not a functional ortholog to *AtMYB36.* **A.** Extended *SlMYB36* phylogeny. Phylogenetic trees generated using protein sequences of several plant species. AmTr: *Amborella trichopoda*, AT: *Arabidopsis thaliana*, Asparagus: *Asparagus officinalis*, Azfi: *Azolla filiculoides*, Bol: *Brassica oleracea*, Carub: *Capsella rubella*, CA: *Capsicum annuum*, Cc: *Coffea canephora*, Cp: *Cucurbita pepo*, DCAR: *Daucus carota*, Gb: *Ginkgo biloba*, HanXRQ: *Helianthus annuus*, MD: *Malus domestica*, Mapoly: *Marchantia polymorpha*, Medtr: *Medicago truncatula*, Migut: *Mimulus guttatus*, GSMUA: *Musa acuminata*, OIT: *Nicotiana attenuata*, GWHPAAYW: *Nymphaea colorata*, LOC_Os: *Oryza sativa japonica*, Peaxi: *Petunia axillaris*, Pp: *Physcomitrella patens*, MA: *Picea abies*, Potri: *Populus trichocarpa*, Semoe: *Selaginella moellendorffii*, Seita: *Setaria italica*, Solyc: *Solanum lycopersicum*, PGSC: *Solanum tuberosum*, Sobic: *Sorghum bicolor*, Thecc: *Theobroma cacao*, VIT: *Vitis vinifera*, Zm: *Zea mays*. **B.** Detailed *MYB36* phylogeny from green square with *AtMYB36 (At5g57620*) highlighted in red, *SlMYB36* (Solyc07g006750) highlighted in pink, and *SlMYB36-b (Solyc04g077260*) highlighted in blue. **C.** Expression patterns of two *SlMYB36* homologs from the *MYB36* phylogeny in individual cell-types from data in Kajala et al. 2021. Legend on the left represents the Log2 maximum value across all conditions and legend on the right represent the Log2(x)/Median. **D.** Cross section of *R. rhizogenes* generated mutant allele *myb36-b hairy root line 8* (*slmyb36-b hr-8*) for *Solyc04g077260*. Left panel: whole root cross section. Right panel magnifies the endodermis. Blue asterisks show the polar lignin cap in the exodermis is not affected. Scale bar = 50 μm.

**Supplemental Figure 6.**
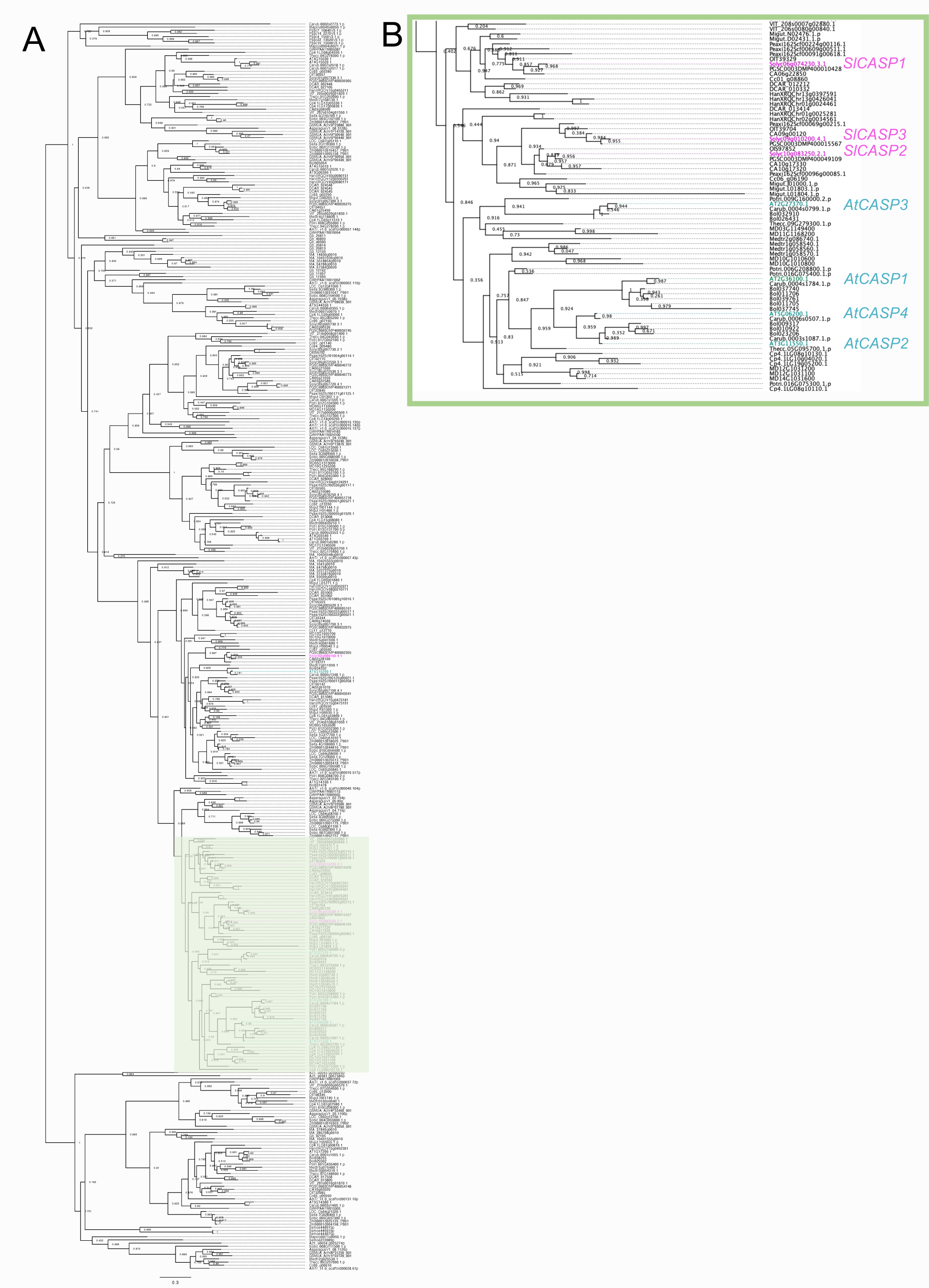
*SlCASP* phylogeny. **A.** Extended *SlCASP* phylogeny. Phylogenetic trees generated using protein sequences of several plant species. AmTr: *Amborella trichopoda*, AT: *Arabidopsis thaliana*, Asparagus: *Asparagus officinalis*, Azfi: *Azolla filiculoides*, Bol: *Brassica oleracea*, Carub: *Capsella rubella*, CA: *Capsicum annuum*, Cc: *Coffea canephora*, Cp: *Cucurbita pepo*, DCAR: *Daucus carota*, Gb: *Ginkgo biloba*, HanXRQ: *Helianthus annuus*, MD: *Malus domestica*, Mapoly: *Marchantia polymorpha*, Medtr: *Medicago truncatula*, Migut: *Mimulus guttatus*, GSMUA: *Musa acuminata*, OIT: *Nicotiana attenuata*, GWHPAAYW: *Nymphaea colorata*, LOC_Os: *Oryza sativa japonica*, Peaxi: *Petunia axillaris*, Pp: *Physcomitrella patens*, MA: *Picea abies*, Potri: *Populus trichocarpa*, Semoe: *Selaginella moellendorffii*, Seita: *Setaria italica*, Solyc: *Solanum lycopersicum*, PGSC: *Solanum tuberosum*, Sobic: *Sorghum bicolor*, Thecc: *Theobroma cacao*, VIT: *Vitis vinifera*, Zm: *Zea mays*. **B.** Detailed phylogeny from green square in A. *AtCASP* and *SlCASP* genes are colored in blue and pink respectively.

**Supplemental Figure 7.**
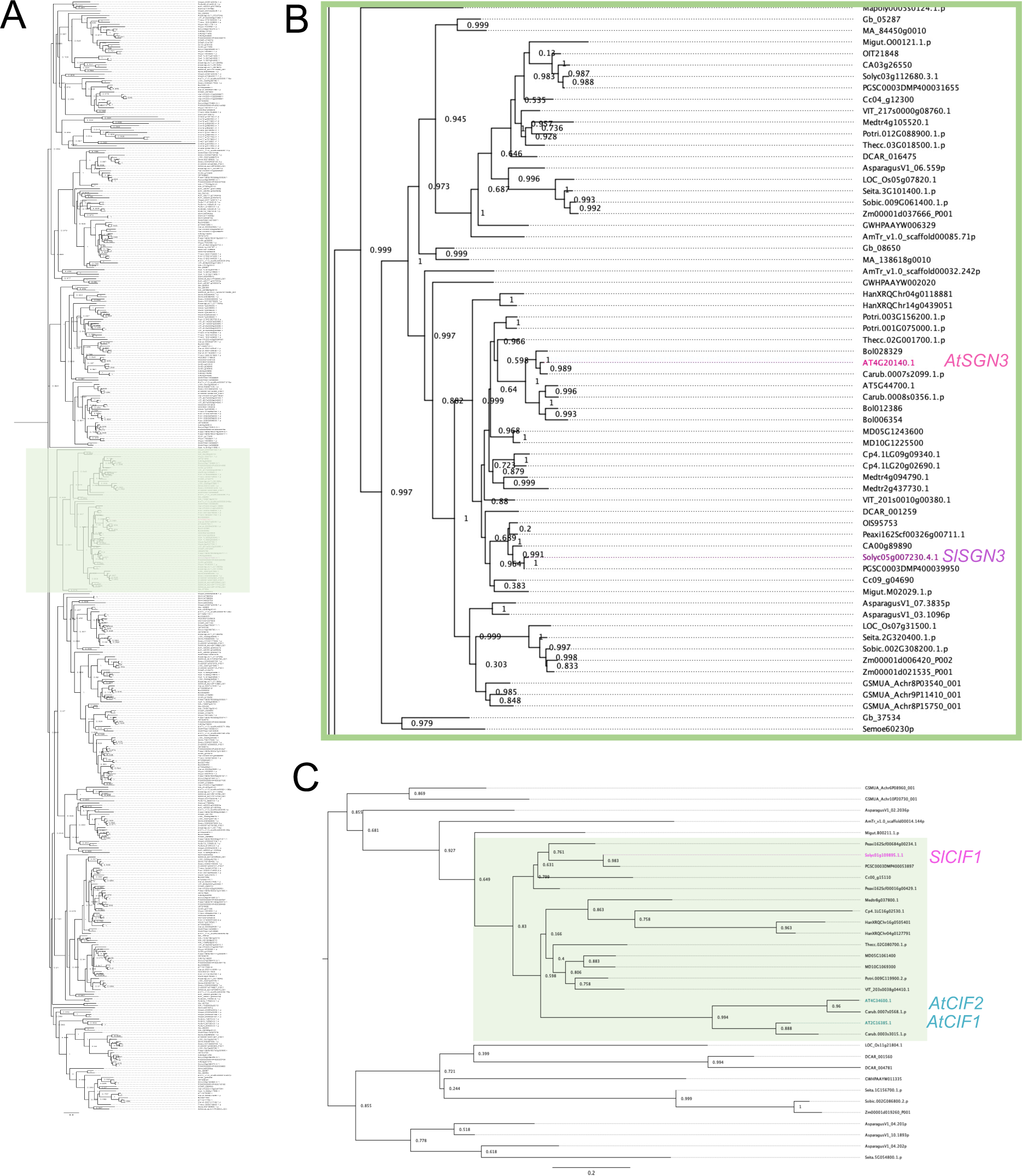
*SlSGN3* and *SlCIF1* phylogeny. **A.** Extended SlSGN3 phylogeny. Phylogenetic trees generated using protein sequences of several plant species. AmTr: *Amborella trichopoda*, AT: *Arabidopsis thaliana*, Asparagus: *Asparagus officinalis*, Azfi: *Azolla filiculoides*, Bol: *Brassica oleracea*, Carub: *Capsella rubella*, CA: *Capsicum annuum*, Cc: *Coffea canephora*, Cp: *Cucurbita pepo*, DCAR: *Daucus carota*, Gb: *Ginkgo biloba*, HanXRQ: *Helianthus annuus*, MD: *Malus domestica*, Mapoly: *Marchantia polymorpha*, Medtr: *Medicago truncatula*, Migut: *Mimulus guttatus*, GSMUA: *Musa acuminata*, OIT: *Nicotiana attenuata*, GWHPAAYW: *Nymphaea colorata*, LOC_Os: *Oryza sativa japonica*, Peaxi: *Petunia axillaris*, Pp: *Physcomitrella patens*, MA: *Picea abies*, Potri: *Populus trichocarpa*, Semoe: *Selaginella moellendorffii*, Seita: *Setaria italica*, Solyc: *Solanum lycopersicum*, PGSC: *Solanum tuberosum*, Sobic: *Sorghum bicolor*, Thecc: *Theobroma cacao*, VIT: *Vitis vinifera*, Zm: *Zea mays*. **B.** Detailed phylogeny from green square in A. AtSGN3 and SlSGN3 genes are colored in pink and purple respectively. **C.** Extended SlCIF1 phylogeny. Phylogenetic trees generated using protein sequences of several plant species. AmTr: *Amborella trichopoda*, AT: *Arabidopsis thaliana*, Asparagus: *Asparagus officinalis*, Azfi: *Azolla filiculoides*, Bol: *Brassica oleracea*, Carub: *Capsella rubella*, CA: *Capsicum annuum*, Cc: *Coffea canephora*, Cp: *Cucurbita pepo*, DCAR: *Daucus carota*, Gb: *Ginkgo biloba*, HanXRQ: *Helianthus annuus*, MD: *Malus domestica*, Mapoly: *Marchantia polymorpha*, Medtr: *Medicago truncatula*, Migut: *Mimulus guttatus*, GSMUA: *Musa acuminata*, OIT: *Nicotiana attenuata*, GWHPAAYW: *Nymphaea colorata*, LOC_Os: *Oryza sativa japonica*, Peaxi: *Petunia axillaris*, Pp: *Physcomitrella patens*, MA: *Picea abies*, Potri: *Populus trichocarpa*, Semoe: *Selaginella moellendorffii*, Seita: *Setaria italica*, Solyc: *Solanum lycopersicum*, PGSC: *Solanum tuberosum*, Sobic: *Sorghum bicolor*, Thecc: *Theobroma cacao*, VIT: *Vitis vinifera*, Zm: *Zea mays*. *AtCIF1* and *AtCIF2* genes are colored in cyan and *SlCIF1* is colored in magenta.

**Supplemental Figure 8.**
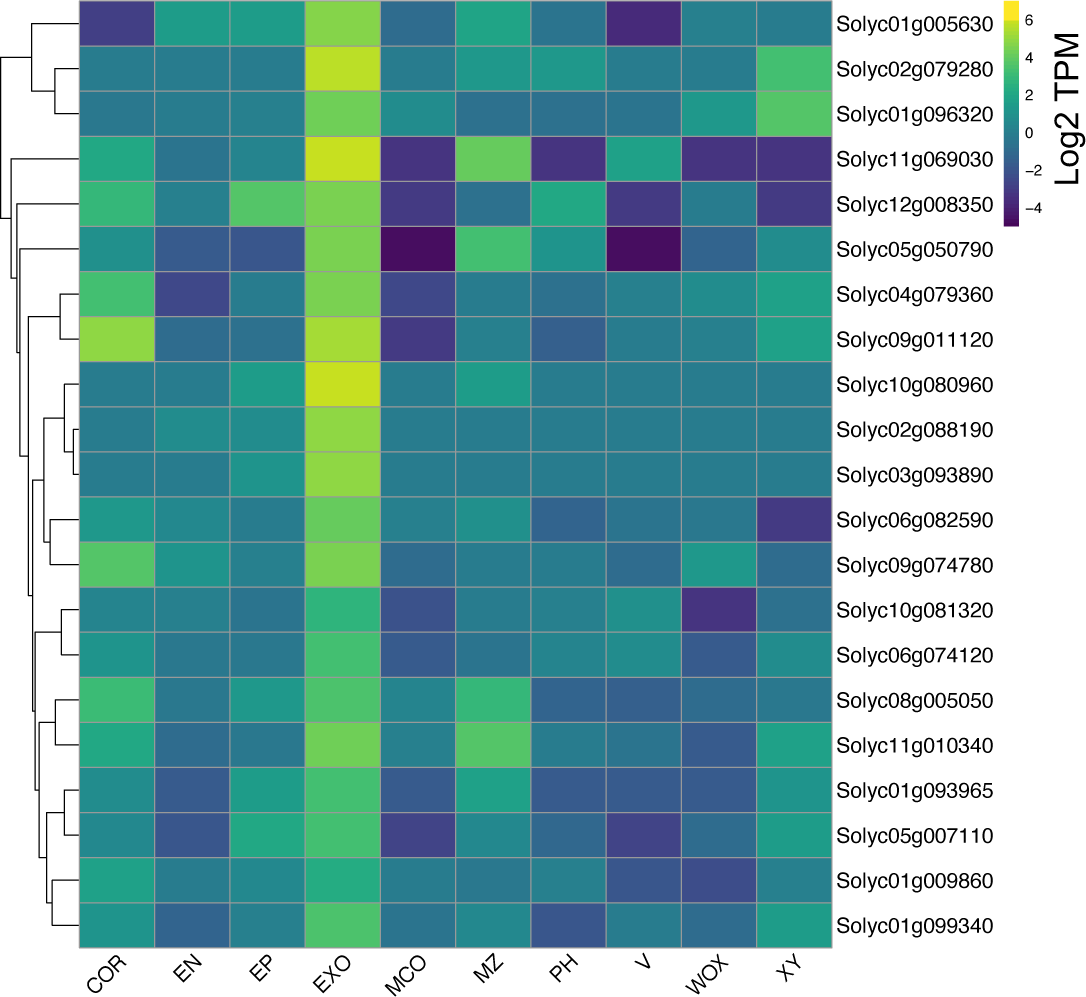
The expression of exodermal-enriched transcription factors from Kajala et al., (2021) selected for a hairy root mutant screen. The column names: Epidermis (EP), Exodermis (EXO), Cortex (COR), EN (Endodermis), MCO (Meristematic cortex), Meristematic zone (MZ), Phloem (PH), Vasculature (V), Quiescent center (WOX5), Xylem (XY).

**Supplemental Figure 9.**
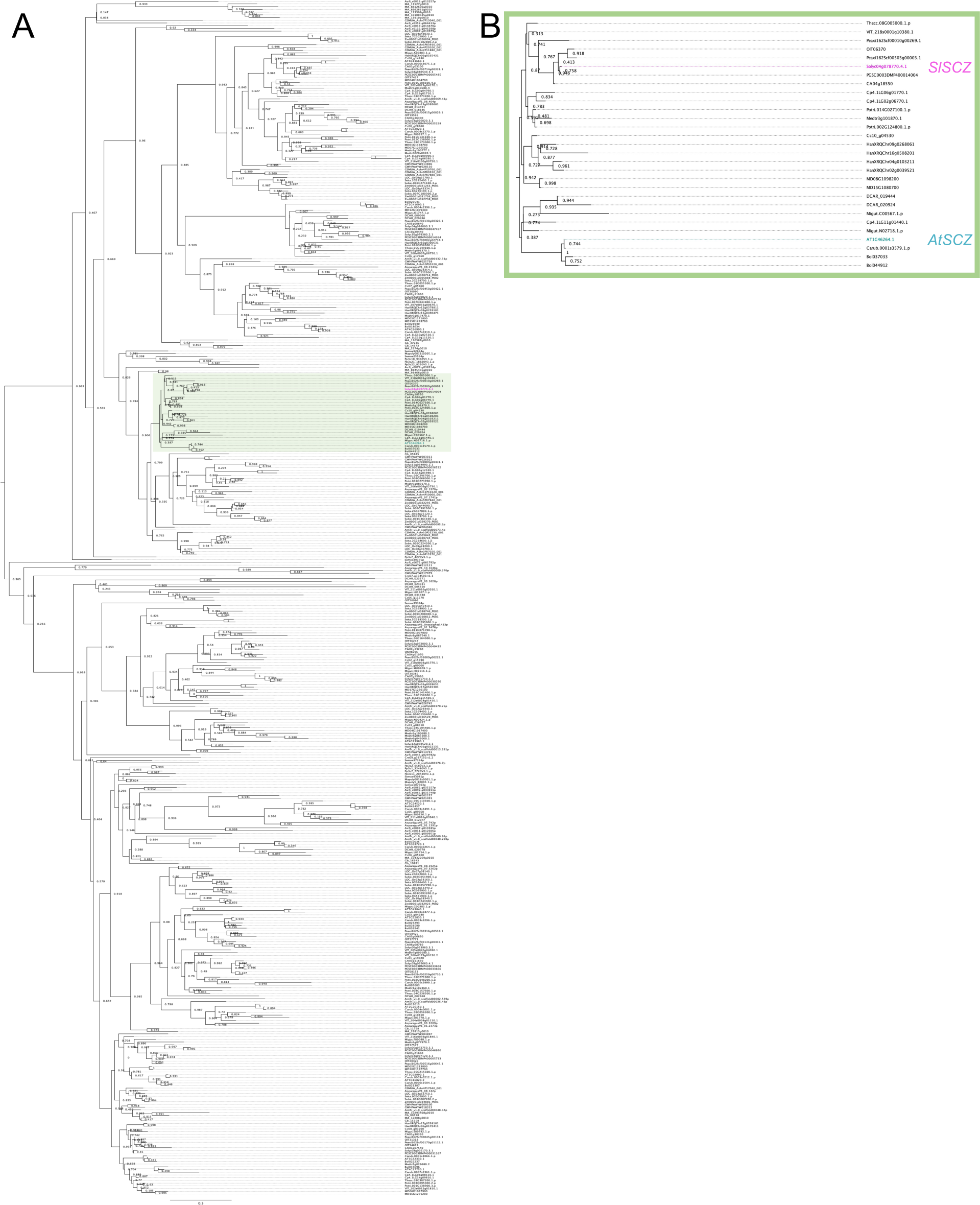
*SlSCZ* phylogeny. **A.** Extended SlSCZ phylogeny. Phylogenetic trees generated using protein sequences of several plant species. AmTr: *Amborella trichopoda*, AT: *Arabidopsis thaliana*, Asparagus: *Asparagus officinalis*, Azfi: *Azolla filiculoides*, Bol: *Brassica oleracea*, Carub: *Capsella rubella*, CA: *Capsicum annuum*, Cc: *Coffea canephora*, Cp: *Cucurbita pepo*, DCAR: *Daucus carota*, Gb: *Ginkgo biloba*, HanXRQ: *Helianthus annuus*, MD: *Malus domestica*, Mapoly: *Marchantia polymorpha*, Medtr: *Medicago truncatula*, Migut: *Mimulus guttatus*, GSMUA: *Musa acuminata*, OIT: *Nicotiana attenuata*, GWHPAAYW: *Nymphaea colorata*, LOC_Os: *Oryza sativa japonica*, Peaxi: *Petunia axillaris*, Pp: *Physcomitrella patens*, MA: *Picea abies*, Potri: *Populus trichocarpa*, Semoe: *Selaginella moellendorffii*, Seita: *Setaria italica*, Solyc: *Solanum lycopersicum*, PGSC: *Solanum tuberosum*, Sobic: *Sorghum bicolor*, Thecc: *Theobroma cacao*, VIT: *Vitis vinifera*, Zm: *Zea mays*. **B.** Detailed phylogeny from the green square in A. AtSCZ and SlSCZ genes are colored in cyan and pink respectively.

**Supplemental Figure 10.**
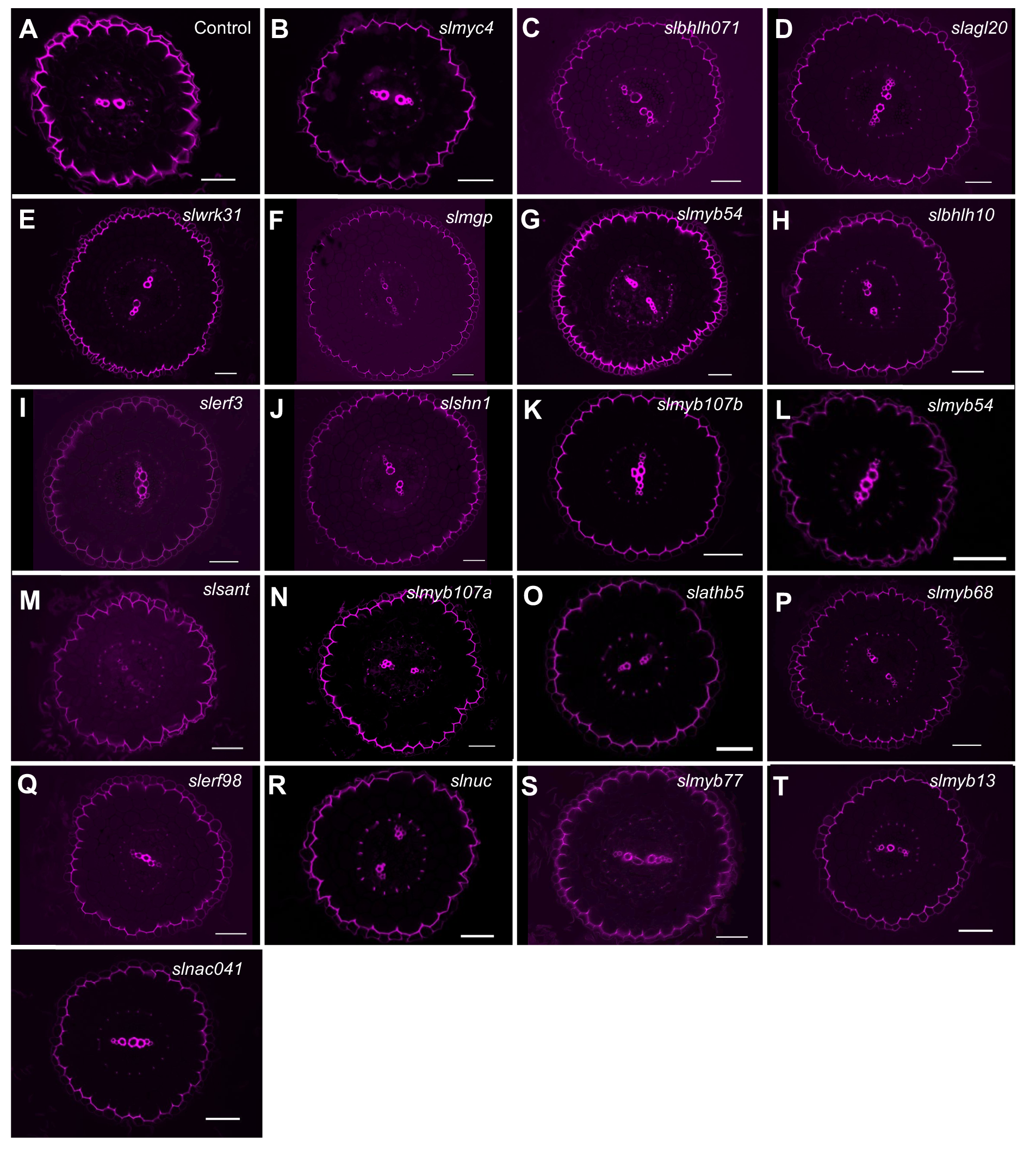
CRISPR-edited hairy root mutants for exodermal-enriched transcription factors. Guide RNAs and edited mutations are found in Supplemental Table 1. For all panels: Root cross sections stained with Basic Fuchsin. **A**. Control = nontransformed hairy root. **B**. *slmyc4* hairy root line 2 (*Solyc08g005050*). **C**. *slbhlh071-1* hairy root line 9 (*Solyc11g010340*). **D**. *slagl20* hairy root line 9 (*Solyc01g093965*). **E**. *slwrk31* hairy root line 3 *(Solyc05g007110*). **F**. *slmgp* hairy root line 17 (*Solyc01g099340*). **G**. *slmyb54* hairy root line 2 (*Solyc10g081320*). **H**. *slblh10* hairy root line 16A (*Solyc06g074120*). **I**. *slerf3* hairy root line 3 *(Solyc06g082590*). **J**. *slshn1 hairy root line 5* (*Solyc01g005630*). **K**. *slmyb107b* hairy root line 6 (Solyc02g088190). **L**. *slmyb54-1* (*Solyc03g093890*). **M**. *slsant* hairy root line 7 (*Solyc10g080960*). **N**. *slmyb107a* hairy root line 15 (*Solyc02g079280*). **O**. *slathb5* hairy root line 8 (*Solyc01g096320*). **P**. *slmyb68* hairy root line 7 *(Solyc11g069030*). **Q**. *slerf98* hairy root line 12 (*Solyc05g050790*). **R**. *slnuc* hairy root line 11 (*Solyc09g074780*). **S**. *slmyb77* hairy root line 6 (*Solyc04g079360*). **T**. *slmyb13* hairy root line 7 (*Solyc08g008480*). **U**. *slnac041* hairy root line 8 (*Solyc01g009860*). Scale bar = 50 μm.

**Supplemental Figure 11.**
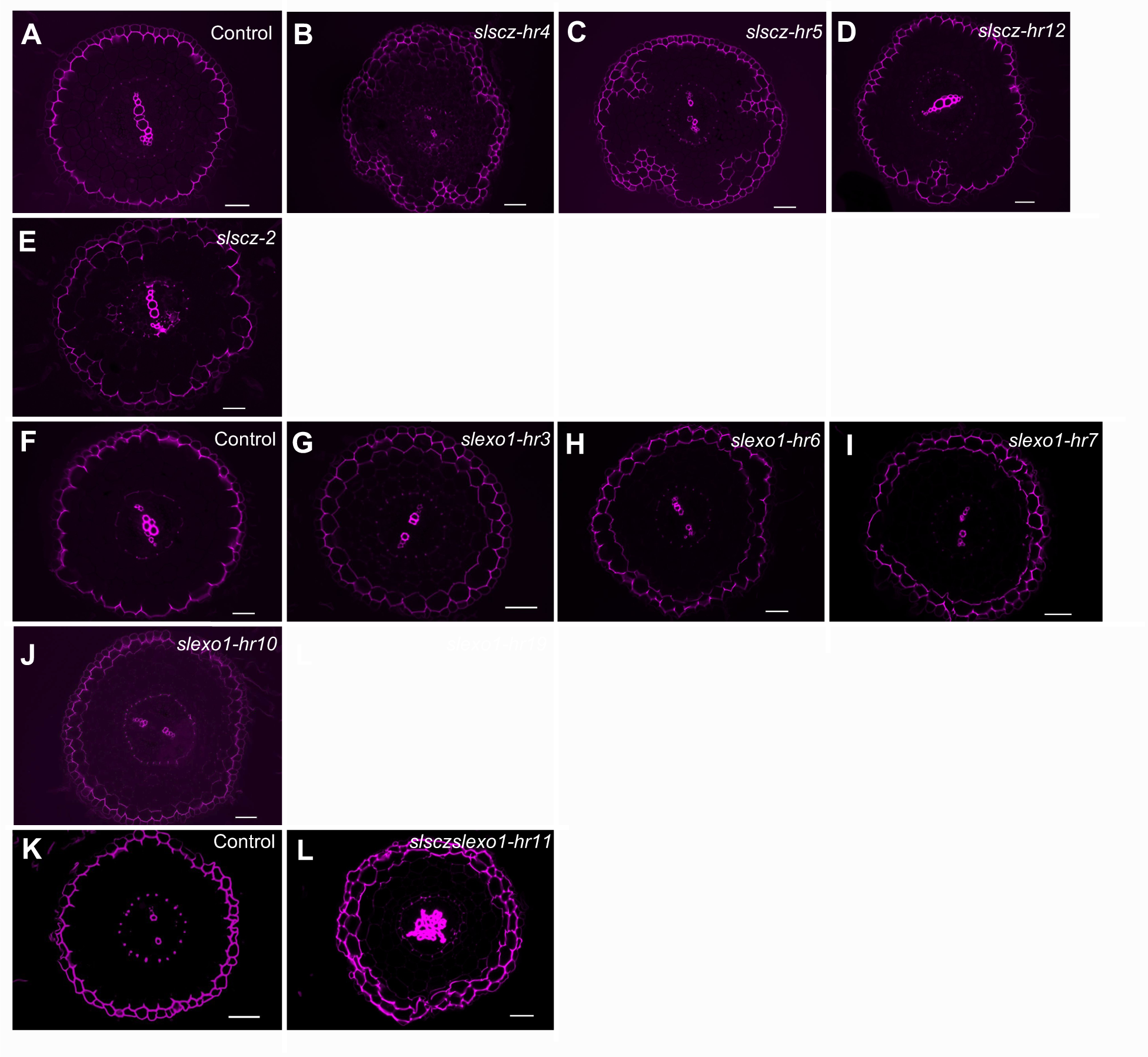
*slscz and slexo1* hairy root mutant lines. **A.** Control cross section stained with fuchsin. **B**. *slscz hairy root line 4* (*slscz-hr4*) cross section stained with fuchsin. **C**. *slscz hairy root line 5* (*slscz-hr5*) cross section stained with fuchsin. **D**. *slscz hairy root line 12* (*slscz-hr12*) cross section stained with fuchsin. **E**. *A. tumefaciens*-transformed *slscz-2* line cross section stained with fuchsin. **F**. Control cross section stained with fuchsin. **G**. *slexo1 hairy root line 3* (*slexo1-hr3*) cross section stained with fuchsin. **H**. *slexo1 hairy root line 6* (*slexo1-hr6*) cross section stained with fuchsin. **I**. *slexo1 hairy root line 7* (*slexo1-hr7*) cross section stained with fuchsin. **J**. *slexo1 hairy root line 10* (*slexo1-hr10*) cross section stained with fuchsin. **K**. Control cross section stained with fuchsin. **L**. *slexoslscz hairy root line 11* (*slsczslexo1-hr11*) double mutant cross section stained with fuchsin. Scale bar = 50 μm.

**Supplemental Figure 12.**
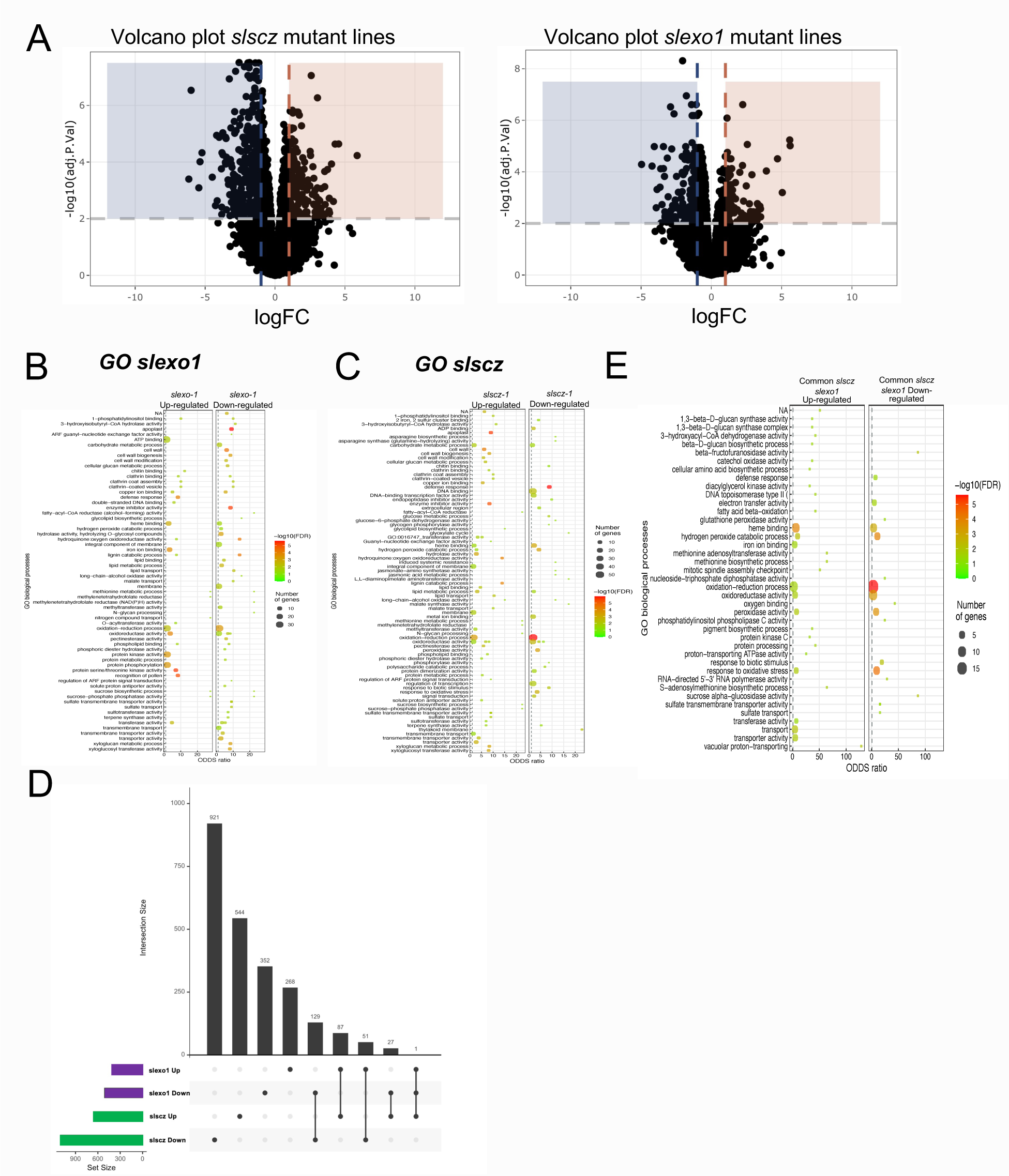
Differentially expressed genes (DEG) in *slexo1* and *slscz* hairy root mutants and associated enriched GO categories. **A**. Volcano plot with the DEG from *slexo1* and *slscz* hairy root mutants. **B**. Enriched GO terms in *slexo1* for upregulated genes relative to wild type and downregulated genes relative to wild type (FC 1.3 p-value <0.05). **C**. Enriched GO terms in *slscz* for upregulated genes relative to wild type and downregulated genes relative to wild type (FC 1.3 p-value <0.05). **D.** UpSet diagram indicating overlap between each of the genotypes for upregulated genes relative to wild type and downregulated genes relative to wild type, with the same FDR and fold-change in C. **E.** GO categories from the common upregulated genes relative to wild type and downregulated genes relative to wild type (FC 1.3 p-value <0.05) from the common *slexo1* and *slscz* genes presented in the Venn diagram from panel B.

**Supplemental Figure 13.**
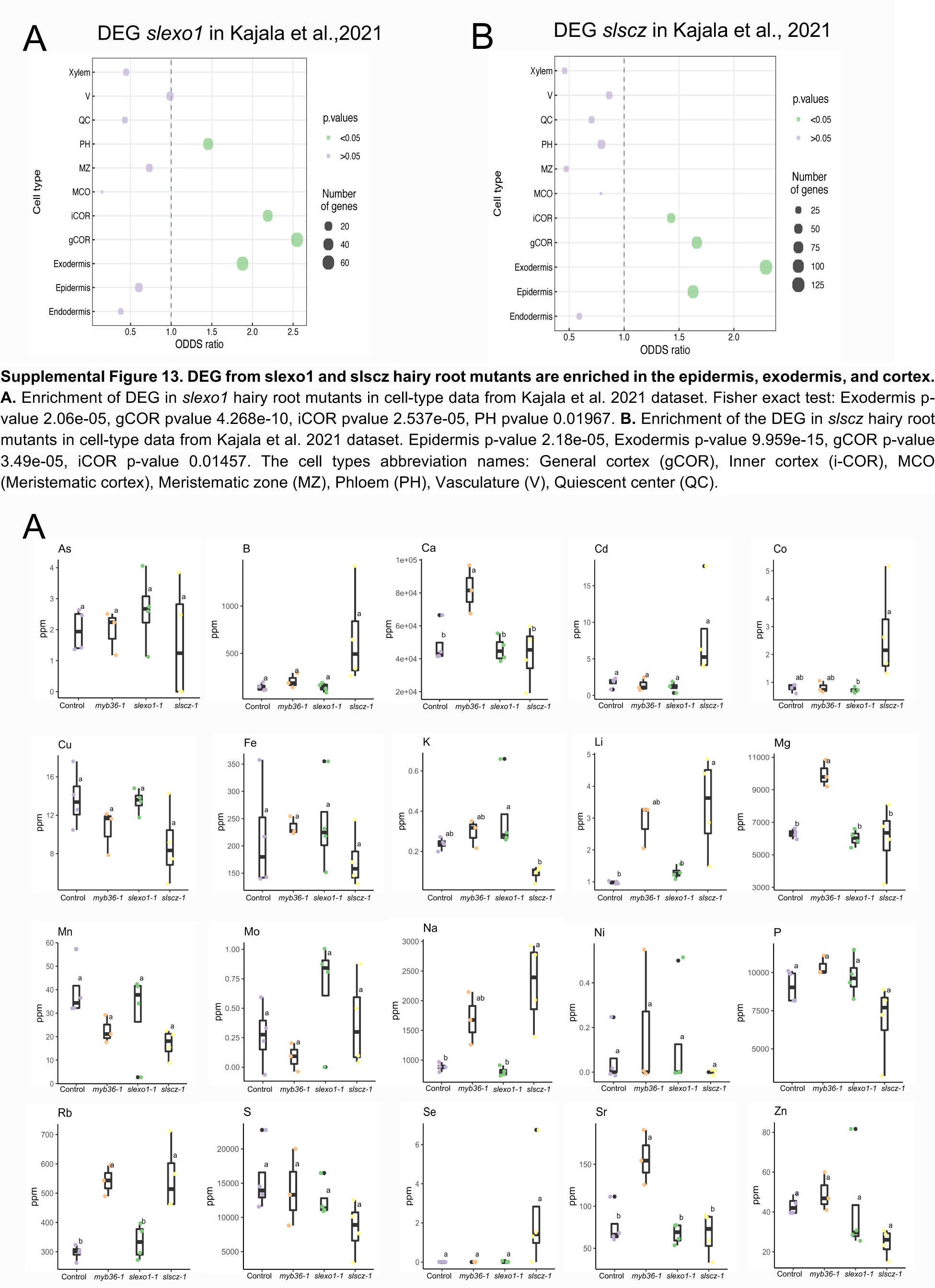
DEG from slexo1 and slscz hairy root mutants are enriched in the epidermis, exodermis, and cortex. **A.** Enrichment of DEG in *slexo1* hairy root mutants in cell-type data from Kajala et al. 2021 dataset. Fisher exact test: Exodermis p-value 2.06e-05, gCOR pvalue 4.268e-10, iCOR pvalue 2.537e-05, PH pvalue 0.01967. **B.** Enrichment of the DEG in *slscz* hairy root mutants in cell-type data from Kajala et al. 2021 dataset. Epidermis p-value 2.18e-05, Exodermis p-value 9.959e-15, gCOR p-value 3.49e-05, iCOR p-value 0.01457. The cell types abbreviation names: General cortex (gCOR), Inner cortex (i-COR), MCO (Meristematic cortex), Meristematic zone (MZ), Phloem (PH), Vasculature (V), Quiescent center (QC).

**Supplemental Figure 14.**
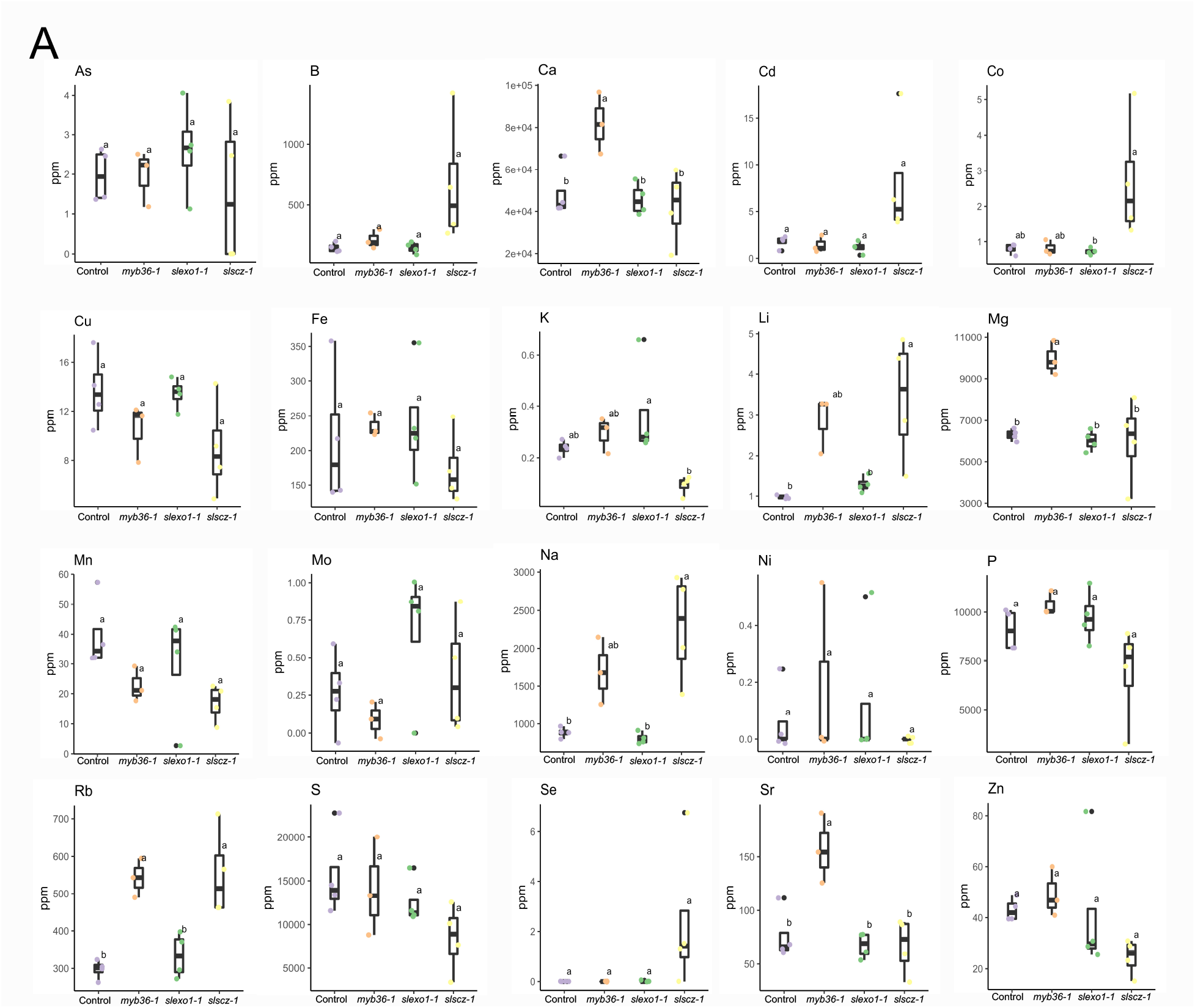
Ion content in *slexo1-1*, *slscz-1*, *slmyb36-*1 and wild-type. **A.** Ion content in ppm (Parts per million) for As (Arsenic), B (Boron), Ca (Calcium), Cd (Cadmium), Co (Cobalt), Cu (Cupper), Fe (Iron), K (Potassium), Li (Lithium), Mg (Magnesium), Mn (Manganese), Mo (Molibdenum), Na (Sodium), Ni (Nikel), P (Phosphorous), Rb (Rubidium), S (Sulfur), Se (Selenium), Sr (Strontium), and Zn (Zinc). Statistical significance was determined by ANOVA with a post-hoc Tukey HSD test.

**Supplemental Table 1.** CRISPR guides and mutation indels description.

**Supplemental Table 2.** Primer sequences.

**Supplemental Table 3.** Primers transcriptional and translational fusions.

## AUTHORS CONTRIBUTION

**Conceptualization**: CM; SMB.

**Methodology**: CM; JCdP; NG; KK (Exodermis TF); NS (Exodermis TF); JBS (Exodermis TF); SMB.

**Investigation**: CM; KWM (Figure 2, Figure 3, Supplemental Figure 5, Supplemental Figure 6, Supplemental Figure 7, Supplemental Figure 9, Supplemental Figure 10, Supplemental Figure 11); LSM (Figure 2, Supplemental Figure 4, Supplemental Figure 8); DdB (Figure 1C-D, Supplemental Figure 3); RU (Figure 1A-C-D, Supplemental Figure 3); SMB.

**Computational investigation**: CM; ACP.

**Formal analyses**: CM; SMB.

**Resources**: CM; KWM (cloning, hairy root transformation, genotyping, CRISPR screen, Phylogenetic analyses, *A.tumefaciens* transgenics); LSM (design CRISPR); ACP (CRISPR cloning); GAM (CRISPR screen); MG (CRISPR cloning).

**Writing - original draft**: CM; SMB.

**Visualization/Figures**: CM; KWM; LSM.

**Supervision**: CM; JCdP; NK; SMB.

**Editing**: All authors contributed to the edition of the manuscript.

## ACKNOWLEDMENTS

We would like to thank A.M. Bagman for experimental assistance; P.A. Merlos de Santa Ana for insightful comments on the project and manuscript. Funding was as follows: CM by MSCA GF 655406. CM, KWM, SMB by NSF 2118017. SMB, ACP, JBS, NS by NSF PGRP IOS-1856749 and IOS-211980. LSM by BARD FI-570-2018 and HFSP #RGP0067. GAM by NSF PRFB IOS-1907008. RU by EMBO Long-term Fellowship ALTF 1046-2015. KK by SKR Postdoctoral Fellowship and MSCA RI Fellowship 790057. JCdP by BIO2017-82209-R. SMB by HHMI 55108506.

